# Tracing colorectal malignancy transformation from cell to tissue scale

**DOI:** 10.1101/2025.06.23.660674

**Authors:** Helena L. Crowell, Irene Ruano, Zedong Hu, Yourae Hong, Gin Caratù, Hubert Piessevaux, Ashley Heck, Rachel Liu, Max Walter, Megan Vandenberg, Kim Young, Dan McGuire, Evelyn Metzger, Margaret L. Hoang, Joseph M. Beechem, Sabine Tejpar, Anna Pascual-Reguant, Holger Heyn

## Abstract

The transformation of normal intestinal epithelium into colorectal cancer (CRC) involves coordinated changes across molecular, cellular, and architectural scales; yet, how these layers integrate remains poorly resolved. Here, we survey colorectal tumorigenesis by combining whole-transcriptome spatial molecular imaging (WTx CosMx SMI) with single-nucleus RNA-sequencing (snPATHO-seq) and digital histopathology on colon samples containing reference mucosa, adenomas and carcinomas, as well as a metastatic lymph node. Leveraging (discrete) histological annotations and (continuous) data-driven trajectories, we quantify the dynamics of cellular density, heterogeneity, function and signaling along the reference-adenoma–carcinoma axis, which is concordant in its spatial and molecular definition. This combination of analytical approaches across different data views let us chart tissue transformation across dimensions (physical/transcriptional) and scales (cell/tissue).

We resolve ∼3.5 million cells into 43 epithelial, immune, and stromal subpopulations that exhibit a bifurcating tumor evolution: On the one hand, LGR5+ stem-like epithelial cells are enriched in highly homogeneous proliferative tumor cores. On the other hand, MMP7+ fetal-like states are restricted to immunosuppressive invasive fronts, rich in cancer-associated fibroblasts (CAFs) and tumor-associated macrophages. These subpopulations form concentric spatial layers that organize transformed regions, and their compound aligns with histological malignancy. We further define ‘transition crypts’ – single colonic crypts with divided histological and transcriptional makeup – that arise from rare crypt fusion or abrupt transformation events. Finally, we trace MMP7+ fetal-like tumor cell states and concomitant myofibroblast-like FAP+ CAFs to lymphovascular invasion sites and matched lymph node metastases, thereby recapitulating invasive programs at single-cell resolution and across sites.

In all, we present single cell- and WTx-resolved spatial data that are among the first of their kind. These open up spatial-centric, out-of-the-box analytical avenues to resolve the molecular, cellular and architectural dynamics that attend tissue transformation during CRC onset, progression and dissemination.

## Introduction

Histopathology has long served as the gold standard for the diagnosis of cancer. Hematoxylin and eosin (H&E) staining enables the visualization of tissue architecture and cellular morphology, and plays a central role in tumor grading, staging, and prognosis^[1,2]^. Digital pathology has further enhanced these capabilities by enabling standardized image analysis and computational modeling^[3]^. Despite these advances, histopathology presents limitations in capturing the full molecular and cellular complexity of cancer needed to identify early steps of transformation and to resolve intra-tumoral heterogeneity^[4,5]^. In contrast, cancer heterogeneity and evolution have been delineated across genomic scales, initially using bulk and, more recently, single-cell sequencing methods that have allowed deciphering clonal evolution and plasticity at the resolution of individual cancer cells.

However, tumor cells do not develop in isolation, but grow in a complex and dynamic microenvironment of stromal and immune cells, which shape the tumor over time and contribute to the clinical progression and treatment response. Numerous single-cell sequencing studies have mapped the landscape of cells in the tumor microenvironment (TME) and generated catalogs of cell types and states across patients and cancer types^[6,7]^. Technologies to project the tumor ecosystem into the spatial context have become available more recently^[8,9,10,11]^. Especially spatial transcriptomics (ST) methods now enable to digitize tumor cells and their microenvironment to map the complex architecture from cell- to tissue-scales. Here, the capture resolution of sequencing-based approaches has increased from 100um (ST^[12]^) to 50um (Visium), 2um (HDST^[13]^) and 250nm (Stereo-seq^[14]^). Meanwhile, imaging-based approaches have improved from resolving 100s of targets^[15]^ to the protein-coding transcriptome^[16]^. The latter now allows a paradigm shift from dissociating cells for single-cell RNA sequencing (scRNA-seq) to directly charting cellular phenotypes in their natural environment. It also overcomes analytical challenges, including library size-based normalization^[17,18]^ and the integration with single-cell reference data, which have been problematic for smaller gene panels. Further, we advance from inference-based interaction analysis of tumor, immune and stromal cells to the *bona fide* detection of co-localization, neighborhoods and, eventually, to derive cellular maps of spatially-resolved tumor ecosystems.

The intestinal epithelium provides an excellent model system where, in health, the transcriptional differentiation trajectory naturally aligns with a physical axis. It is a rapidly renewing tissue that maintains homeostasis through a tightly regulated balance of cell proliferation and differentiation^[19]^. This continuous turnover, essential for the long-term function of the gastrointestinal tract, occurs in a spatially orchestrated manner along the crypt axis (from the basal columnar stem cells towards the luminal interface), and is reinforced by cellular interactions and the sensing of local microenvironmental factors^[20,21]^. Colorectal carcinoma (CRC) originates from colonic crypt epithelial cells that transform from normal to malignant cancer cells through well-characterized processes involving intermediate premalignant polyp stages (adenomas), considered as major precancerous lesions^[22]^. Today, we have a comprehensive view of the adenoma-carcinoma sequence from the genomic, transcriptomic and proteomic perspective^[23,24,25,26]^, and at single-cell scale^[27,28,29]^. Yet, the cell intrinsic and extrinsic dependencies of premalignant neoplastic transformation and, in particular, their spatial orchestration remain elusive.

In this study, we apply whole-transcriptome (WTx) spatial molecular imaging (CosMx SMI^®^, Bruker) combined with single-nucleus RNA sequencing (snPATHO-seq) and digital histopathology to analyze normal, premalignant and primary CRC samples and matched lymph node metastases. Importantly, the specimens contained healthy reference mucosa, tubulovillous adenomas (TVA; premalignant polyp lesions) and tumor regions (CRC) within the same section. This allowed us to model tumor evolution at single-cell resolution by ordering the epithelial transformation process along a pseudotime axis. Projected into their spatial coordinates, we followed the tumor, stromal and immune cell co-evolution across spatial scales that represent the interaction of individual cells, co-localization niches, functional units (including crypts and vessels), and tumor-wide architectures. This work combines concepts from classical histopathology and single-cell omics with out-of-the-box spatial analyses, made possible through transcriptome-wide, single cell-resolved spatial molecular imaging, to map the cellular ecosystem along CRC evolution and its architectural traits.

## Results

### Combined CosMx WTx and digital histopathology resolves heterogeneity across and within tissue domains

We acquired WTx CosMx SMI data of eight colon sections and one draining lymph node from seven CRC patients, imaging 1,638 fields of view (FOV). We captured a total scanning area of ∼4.3cm^2^, calling 3,555,234 cells (Table S1). The CosMx data included 18,878 RNA targets, 50 negative probes (to quantify nonspecific hybridization of the ISH probe) and 2,745 false codes (to quantify misidentification of the reporter readout). It also contained immunofluorescence (IF) markers for B2M/CD298 (membrane), DAPI (nuclei), CD45 (immune), CD68 (macrophages) and PanCK (epithelia); see Fig. S1 and Fig. 1a. Cells with low RNA target counts (in total and per area), exceedingly high non-target counts (negative probes and false codes), or localized very near to FOV borders were removed, retaining ∼3.4M high-quality cells for downstream analysis; see Fig. S4 and Methods.

**Figure 1.**
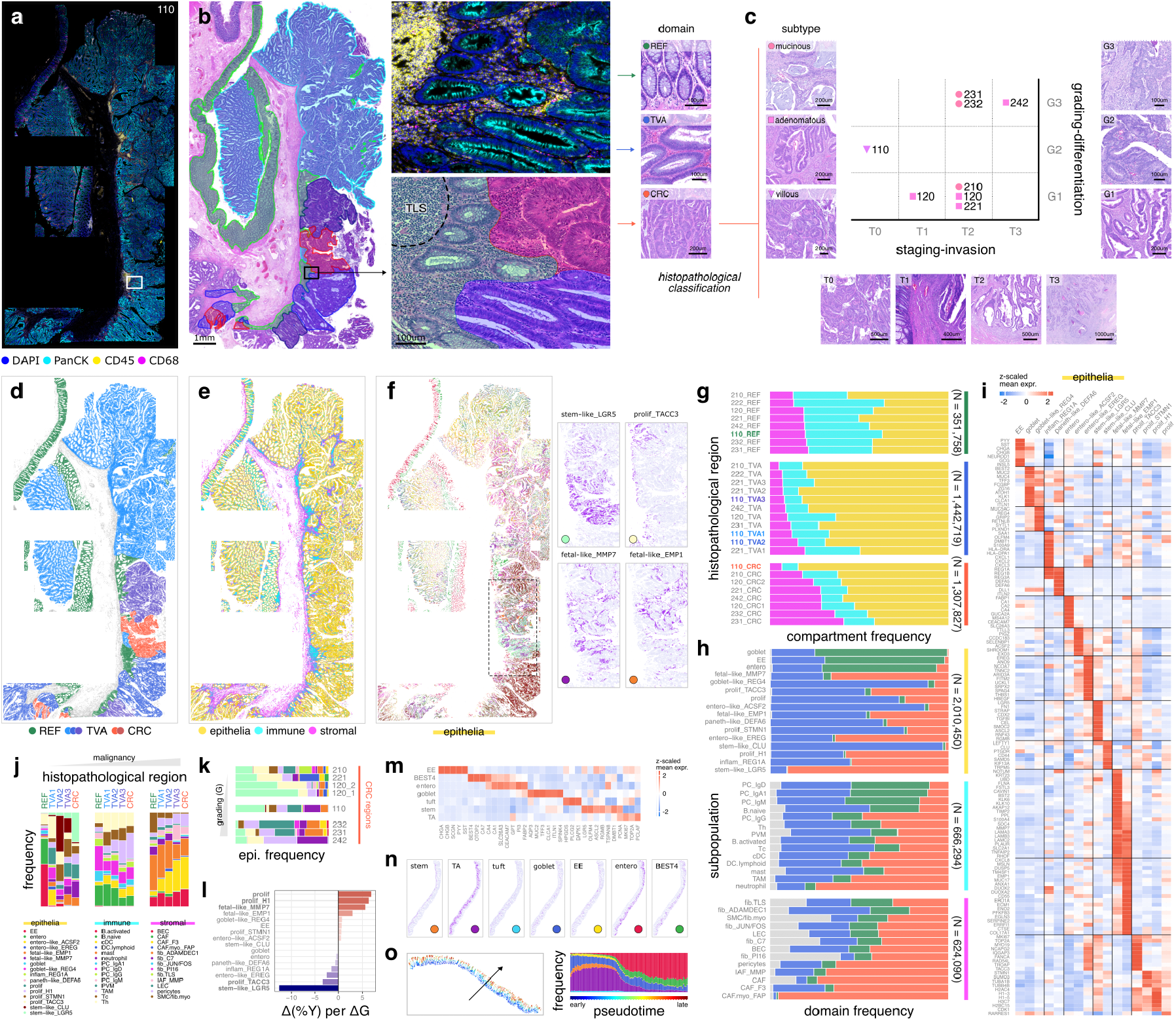
Histopathological and transcriptional tissue characterization. **(a)** Immunofluorescence (IF) composite image (section 110). **(b)** H&E stain overlaid with histopathological annotation of referencelike mucosa (REF), premalignant tubulovillous adenoma (TVA), and colorectal cancer (CRC). Zoomed in region in (a-b) comprises tissue of all types, and a tertiary lymphoid structure (TLS). **(c)** Schematic illustrating the histopathological classification of CRCs in terms of malignancy score, including cell grading/differentiation (well-G1; moderately-G2; or badly differentiated-G3) and staging/invasion (mucosa-T0; submucosa-T1; muscular-T2; serosa-T3). CRCs were also classified according to their growth pattern: mucinous, adenomatous and villous. Representative H&E stains of relevant features are included. **(d-f)** Spatial plots with cells colored by (fltr) histopathological region, compartment, and epithelial sub-population; and, spatial subplots highlighting selected subpopulations individually. **(g)** Frequency of compartments across histopathological regions (gray = unassigned); y-axes ordered by stromal abundance. **(h)** Frequency of histopathological regions across (fttb) epithelial, immune, and stromal sub-populations; y-axes ordered by REF abundance. **(i)** Selected marker genes for epithelial subpopulations (data are across cells from all sections). **(j)** Frequency of subpopulations across section 110’s histopathological regions; x-axes ordered by malignancy. **(k)** Frequency of epithelial subpopulations across CRC regions from all sections. **(l)** Average change in epithelial subpopulation frequencies per histological grade (G). **(m)** Selected marker genes for normal epithelial subpopulations (data are across cells from all sections). **(n)** Spatial plots highlighting normal epithelial subpopulation individually (subset of section 110’s REF domain). **(o)** Spatial plot with cells colored by pseudotime, and frequency of normal epithelial subpopulations across pseudotime (section 110).

Adjacent tissue sections were stained with H&E and used to classify the colon mucosa into three domains: *reference* (REF), *premalignant* tubulovillous adenoma (TVA), and *malignant* colorectal carcinoma (CRC); see Fig. 1b and Methods. Samples were annotated by two pathologists in an independent double-blind protocol. Carcinomas were scored according to their malignancy features at the histopathological level, including cell grading/differentiation (well-G1; moderately-G2; or poorly differentiated-G3) and staging/invasion (mucosa-T0; submucosa-T1; muscular-T2; serosa-T3); see Fig. 1c. This classification was combined with well-defined subtypes of mucinous and adenomatous CRC. Two donors stood out for also presenting villous growth (110) or adenomatous CRC with neuroendocrine differentiation (NED; 242), the most malignant carcinoma in this study. Villous-like carcinoma in donor 110 was only identified as such by one of the pathologists. The neoplastic domains were further stratified into 2-5 regions per section, where various adenomas and/or carcinomas could be histologically distinguished in terms of growth pattern or malignancy features; see Fig. 1b,d, Fig. S2 and Table S3.

To obtain a single-cell reference for cellular annotation *in situ*, we generated snPATHO-seq^[30]^ data for 125,890 cells from adjacent tissue sections (Table S2), which were annotated into 16 broad subpopulations. A set of subpopulation markers were then used to compute pseudobulk profiles for supervised clustering of the CosMx SMI data (using *InSituType*^[31]^; see Methods for details). Nearly 60% of cells (∼2M) were assigned to epithelial populations (Fig. S5a), followed by fibroblasts (∼440k), plasma, T (∼200k each) and myeloid cells (∼180k). The remaining subpopulations comprised 6,7k mast, 60k B, and 100k smooth muscle cells. Subpopulation frequencies were comparable, though variable, across sections (Fig. S5b), and exhibited clear transcriptional profiles that align with canonical gene expression signatures (Fig. S5d). The main drivers of variability were plasma and B cells, contrasted by epithelia, and stromal subpopulations (Fig. S5c and Fig. S6a,b). Consequently, a clear spatial organization of the tissues was already apparent at high-level cell annotation (Fig. S5e).

Next, we grouped subpopulations into three broad biological compartments, namely, epithelia, immune, and stroma (Fig. 1e and Fig. S7b) and subclustered for downstream analysis (Fig. 1f-j). Across all sections, ∼40-60% of cells were epithelia, with immune and stromal cells contributing similar fractions of non-epithelial cells (Fig. S7c). As expected, the number of total counts and uniquely detected features per area was notably higher in epithelia, which were also highest in PanMem and PanCK staining, while immune cells were highest in CD45 and CD68 fluorescence intensities (Fig. 1a,b and Fig. S7d). REF mucosa regions were richer in immune cells (Fig. 1g), consistent with an expanded lamina propria in these areas (Fig. S3). Hyperproliferation of transformed cells resulted in an increased epithelial compartment in TVA and CRC regions (Fig. S2), while the stromal compartment increased in the CRC domain (Fig. S7h), especially in deeply invasive samples (T2 and T3; Fig. 1c) and/or showing fibrous desmoplasia (both detailed in Table S3). Samples with superficial CRC proliferation without ECM remodeling and invasion, such as 110, also showed a decreased stromal compartment in tumor regions (Fig. S7k).

Compartment-wise subclustering yielded 16 epithelial, 14 immune, and 13 stromal subpopulations (Fig. S10). Within the epithelial compartment, we identified three clusters that mostly aligned with reference-like mucosa, both spatially (Fig. 1h and Fig. S10c) and transcriptionally (Fig. 1i). These clusters contained a mixture of the main cell types known to compose healthy colonic crypts. Mapping to (pre)malignant lesions, we identified four highly proliferative subpopulations (prolif, prolif TACC3, prolif STMN1 and prolif H1), three transformed clusters resembling terminally differentiated reference-like states (goblet-like REG4, entero-like ACSF2 and entero-like EREG), a paneth-like population (paneth-like DEFA6), an inflammation-related cluster (inflam REG1A), two partly overlapping fetal states (fetal-like MMP7 and fetal-like EMP1), and two distinct stemness states (stem-like CLU and stem-like LGR5); see Fig. 1h-j. Non-reference epithelia also exhibited distinct spatial patterns (Fig. 1f,j and Fig. S10c,d). In line with being exclusively localized to the tumor core, the stem-like LGR5 population represents cancer stem cells that self-renew and differentiate to other tumor states, sustaining tumor growth in a crypt-like niche^[32,33,34]^. The stem-like CLU cluster represents a quiescent revival stem cell population that emerges upon intestinal injury and regenerates LGR5+ crypt stem cells in a YAP1-dependent manner^[35,34,36,37,38]^. We found this population largely restricted to adenoma regions, together with goblet-like REG4 and entero-like ACSF2. Mapping to both TVA and CRC areas, prolif TACC3 showed evident basal location in the lower crypt epithelium across domains, while fetal-like EMP1 showed a more patchy distribution in CRC, but outlined the apical side of TVAs. The fetal-like MMP7 population outlined the tumor mass in a spatial pattern suggestive of the invasive front, in accordance with previous studies showing that matrix metalloproteinases -MMPs-(particularly MMP7) facilitate invasion by degrading peritumoral extracellular matrix and junctional proteins^[39]^.

We distinguished three types of CRCs that differed in terms of their histopathological malignancy characterization (Fig. 1c and Table S3) and epithelial composition (Fig. 1k). One type was mainly composed of stem-like LGR5 and proli TACC3, and maps to well-differentiated tumors with lowest grading (G1; sections 120, 210 and 221). The second type was composed of fetal-like MMP7 together with fetal-like EMP1, prolif and prolif H1, and maps to poorly-differentiated tumors with G3 grading (sections 231, 232 and 242). The third was a mixture of these types and mapped to section 110, which exhibits a villous-like growth pattern and moderate cell differentiation (G2). Notably, the annotation of this region in particular was histologically ambiguous, but clear on a transcriptional level. Quantitatively, each step-wise advance in grading/differentiation (G) – i.e., increase in malignancy – corresponded to a 5% increase in fetal-like MMP7, prolif and prolif H1 populations, and a 10% decrease in stemlike LGR5 on average (Fig. 1l).

In the immune compartment, we identified four subpopulations of plasma cells expressing different immunoglobulins (PC IgD, PC IgA1, PC IgG and PC IgM), activated B cells (B.activate), naive B cells (B.naive), T helper cells (Th), cytotoxic T cells (Tc), mast cells, and five myeloid subpopulations, including neutrophils, conventional dendritic cells (cDC), lymphoid DCs (DC.lymphoid), tumor-associated macrophages (TAM), and peri-vascular macrophages (PVM) (Fig. S10a and Fig. S11a). Generally, myeloid, B and T cells were predominantly found in adenomas and carcinomas, while PCs were very abundant in REF regions. However, IgG+ PCs were preferentially located in tumor lesions, together with DCs, TAMs and neutrophils (Fig. 1h,j and Fig. S10b,c). The latter were particularly abundant in section 120, and correlated with pathological findings of large necrotic areas with fibrin, viable and degenerated neutrophils, and vasculitis, suggesting an inflammatory reaction surrounding tissue damage (Fig. S10b and Table S3). Interestingly, neutrophils and Tc cells were abundant in less and intermediate malignant samples, but reduced in the most malignant specimens (Fig. 1c and Fig. S10c).

Within the stromal compartment, we found blood (BEC) and lymphatic (LEC) endothelia, smooth muscle cells (SMC/fib.myo) and various subtypes of fibroblasts (fib ADAMDEC1, fib C7, fib PI16, fib.TLS and fib JUN/FOS; the latter representing a population marked by genes related to cell stress, immunomodulation and fibrosis), all distributed across the three domains. Additionally, we identified pericytes, a subset of inflammatory-associated fibroblasts (IAF MMP), and three distinct cancer-associated fibroblast subtypes (CAF, CAF F3 and CAF.myo FAP) that were largely restricted to the tumor domain (Fig. 1h,j, Fig. S10c and Fig. S11b). The CAF.myo FAP population exhibited markers related to a matrix-producing contractile phenotype, several collagen genes, FN1, CTHRC1 and FAP. They also expressed CTHRC1, a proteoglycan implicated in the progression and therapy-resistance of several cancers^[40,41,42]^ and known to promote EMT, angiogenesis, metabolic alterations and ECM formation, primarily through the regulation of pathways like WNT/PCP, TGFB/SMAD, and MEK/ERK^[43]^.

To fully resolve normal epithelial differentiation states, we subclustered cells annotated as REF mucosa (based on histopathology) using Human Cell Atlas colon data as reference^[44]^; see Methods. The resulting subpopulations (intestinal stem cells, transit amplifying -TA-, tuft, enteroendocrine -EE-, goblet and (BEST4+) enterocytes) expressed canonical marker genes (Fig. 1m). They also mapped to their well-characterized positioning along the basal (stem) to apical (BEST4+ enterocytes) crypt axis (Fig. 1n). Transcription-based trajectory inference on these cells (using *slingshot* ^[45]^) yielded a pseudotime ordering that aligned with the positioning of epithelial differentiation states along the same spatial axis (early: basal; late: apical; Fig. 1o).

### Shared architectural and functional changes attend intra-and inter-domain transformations

To further characterize the tissues beyond cellular composition and in light of a recent study demonstrating that increased crypt density augments adenoma formation^[46]^, we quantified cellular densities across histopathological regions (Fig. 2a-b, Fig. S8 and Fig. S9) and epithelial subpopulations (Fig. 2c). Epithelial density correlated not only with histopathology annotations, but also with transcriptional clustering, increasing from normal to transformed epithelial states. While the density of fetal-like states was similar to REF epithelia, stem-like and metaplastic (paneth-like) populations showed the highest density. In contrast, immune cell density decreased from REF to TVA and cancer, being negatively correlated with epithelial transformation/malignancy. Nevertheless, patches of high immune cell density near the muscularis mucosa within the TVA and CRC domains represented tertiary lymphoid structures (TLS). Accordingly, these were enriched with B.naive, Th and DC.lymphoid cells co-localizing with fib.TLS (Fig. S10m), a stromal population characterized by the expression of cytokines known to organize TLS, such as LTB, CCL19 and CXCL13 (Fig. S11b). We further quantified the purity of subpopulations in the radial neighborhood of each cell using entropy as a metric for homogeneity/heterogeneity (low/high entropy) of local microenvironments (see Methods). Cellular entropies decreased from REF to TVA and CRC (Fig. 2d), pointing to tumoral neighborhoods being the most homogeneous. In this case, however, distinct stem-(LGR5) and fetal-like (EMP1) subpopulations had the lowest entropy (Fig. 2e), indicating mostly homophylic interactions. Thus, albeit related, density and homogeneity represent orthogonal phenomena to chart tissue architecture.

**Figure 2.**
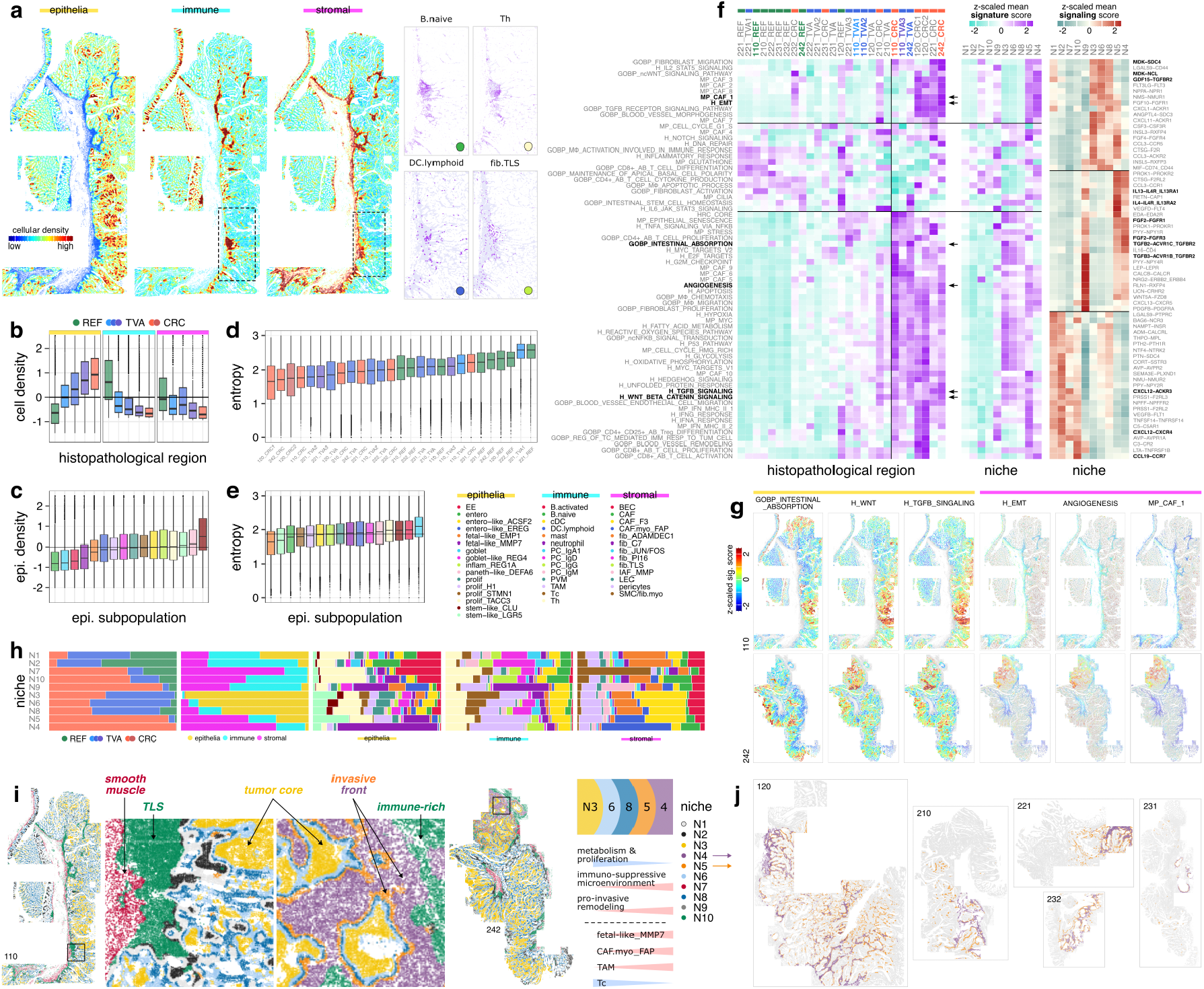
Niche analysis and cell-cell communication. **(a)** Spatial plots colored by (fltr) epithelial, immune and stromal cell density, alongside a region of interest (c.f. Fig. 1a,b,f) highlighting selected subpopulations individually. **(b)** Density (= number of cells within a 50um radius from a given cell, z-scaled across the tissue) of epithelial, immune, and stromal cells across section 110’s histopathological regions; x-axis ordered by malignancy. **(c)** Density of epithelial subpopulations (data across all sections); x-axis ordered by median. **(d-e)** Shannon entropy of subpopulation frequencies in each cell’s 50um radial neighborhoods across histopathological regions and epithelial subpopulations, respectively (data across all section). **(f)** Signature gene set scores across histopathological regions and niches; and, top ranked cell-cell communication interactions across niches (data across all sections). **(g)** Spatial plots colored by selected gene set signature scores, highlighting only epithelial and stromal cells, respectively. **(h)** Frequency of histopathological regions, compartments, epithelial, immune and stromal subpopulations across niches. **(i)** Spatial plots with cells colored by niche (section 110 and 242; c.f. Fig. S12); included is a schematic summarizing niche layering and the accompanying key functional and compositional changes from tumor core to invasive front. **(j)** Spatial plots highlighting cells of niches N4 and N5, marking the invasive front across sections with CRC domain.

We additionally scored signature gene sets to facilitate functional characterization, including key hallmark (e.g., epithelial-mesenchymal transition -EMT-) and malignant transcriptional programs^[6]^ (e.g., CAF); see Fig. 2f,g. Across sections, REF regions were rich in steady-state structural and functional pathways of the lamina propia (maintenance of apical basal cell polarity, cilia, intestinal stem cell homeostasis, glutathione metabolism and fibroblast activation) and homeostatic immune responses (macrophage activation and apoptosis, and T cell differentiation and cytokine production). Major known pathways upregulated during carcinogenesis, such as TGFB, WNT, MYC, NOTCH and Hedgehog, mapped to CRC regions. Similarly, increased tissue damage response (e.g., unfolded protein response, reactive oxygen species -ROS-, DNA repair and p53) and metabolism (hypoxia, oxidative phosphorylation and glycolysis) mapped to these regions, together with pathways related to TME remodeling (CAF, EMT and interferon -IFN-responses). Of note, in section 120, both in the REF and the two distinct CRC regions, we detected an upregulation of IFN pathways, together with pathways related to T cell activity. This aligns with our orthogonal histopathology assessment, showing a high inflammatory state with vasculitis and necrotic regions (Table S3). Additionally, these CRCs presented an enrichment of blood vessel remodeling pathway, which is in line with the activation and remodeling of mucosal micro-vessels occurring in an inflammatory context^[47]^. On the other hand, the CRC in section 242 showed higher angiogenesis and blood vessel morphogenesis pathways, consistent with the need for increased nutrient supply of more aggressive tumors.

To identify concordant microenvironments across sections and domains, we further clustered cells based on the subpopulation composition of their local neighborhood to obtain 10 spatial niches (N1-10; Fig. 2f-j and Fig. S12). Epithelia showed drastic compositional differences across domains (c.f. Fig. 1j and Fig. S10c). When quantifying cellular neighborhoods, we disregarded the identity of epithelial subpopulations (i.e., treating all epithelia as equal). At this, niches only capture shifts in the immune and stromal compartment, as well as epithelial density (see Methods).

The REF domain largely mapped to N1 and N2 that comprised mostly PCs, normal epithelial subpopulations (EE, entero, goblet), and fib ADAMDEC1. In TVA and CRC regions, niche analysis revealed a well-defined spatially layered organization consistent across all samples (Fig. S12). This shared architecture was organized in concentric layers, beginning with a central core dominated by N3 and N6, composed strictly of epithelia, and presenting a mixture of stem-like and proliferative epithelial states (Fig. 2h,i). Moving from the core, N8 was characterized by a gradual transition toward a more proliferative and less stem-like profile. N5 and N4 delineated the invasive front (Fig. 2j), located exclusively in peripheral CRC regions that invaded deeper intestinal layers (represented by N7 and N10). N5 showed an increase in stromal components and fetal-like MMP7 epithelia. Concurrently, there was a coordinated shift in the TME landscape with decreasing Tc cells, and increasing TAMs and invasive CAF.myo FAP populations. Spatially, N4 represented the last layer, with a niche dominated by CAF.myo FAP cells. An exceptions was sample 110, which was graded as a non-invasive carcinoma (T0) and, accordingly, did not have invasive front-related niches. In addition to the layered niche patterns, we captured an inflammatory niche (N9) that was almost entirely confined to CRC regions, and arranged in neutrophil- and IAF MMP-rich patches.

Signature scoring and cell-cell communication analysis (see Methods) across niches functionally delineated the aforementioned compositional and spatial differences. Core epithelial niches in both TVA and CRC regions (N3, N6 and N8) were rich in metabolic and proliferative pathways (glycolysis, fatty acid metabolism, cell cycle, MYC, E2F, p53 or NFKB), and showed an upregulation of mitogenic signaling and growth factors (FGFs, MDK, FLT3LG), angiogenesis, and immune modulation (GDF15, ANGPTL4), pointing to tumor-specific progrowth. The invasive front niches (N5 and N4), in turn, showed upregulation of pathways related to TME remodeling (fibroblast migration, CAFs, angiogenesis, and EMT) and enriched interactions that point to type 2 immune responses (IL-4/IL-13 axis), ECM remodeling (TGFBs, FGF2), macrophage and endothelial support (MIF, VEGFD), and hormone signaling (PYY, LEP, RLN1), all suggestive of a tumor-stroma co-evolution in an immunosuppressive microenvironment. Remaining niches (N1, 2, 7 and 10) showed rather homeostatic and immune surveillance interactions involving neuropeptides and hormones (AVP–AVPRs, PYY–NPYRs, NMU–NMUR2), as well as signals for complement and lymphocyte recruitment (C3–CR2, C5–C5AR1, CXCL12–CXCR4/ACKR3, CCL19–CCR7).

### Epithelial trajectory captures tumor and TME co-evolution from tissue to single-crypt scale

To help characterizing TVA emergence and progression to CRC, we performed transcriptionbased trajectory inference (using *slingshot* ^[45]^; see Methods) of epithelial cells at the tissue-wide scale (Fig. 3a, Fig. S6c,d and Fig. S13a). Hereby, we capture dynamic changes in cellular states from early to late phases, modeling the evolution from REF mucosa through various adenomatous stages to malignant epithelium; considering how non-epithelial cells (physically) organize along this transformation let us further investigate the accompanying immune/stromal adaptations.

**Figure 3.**
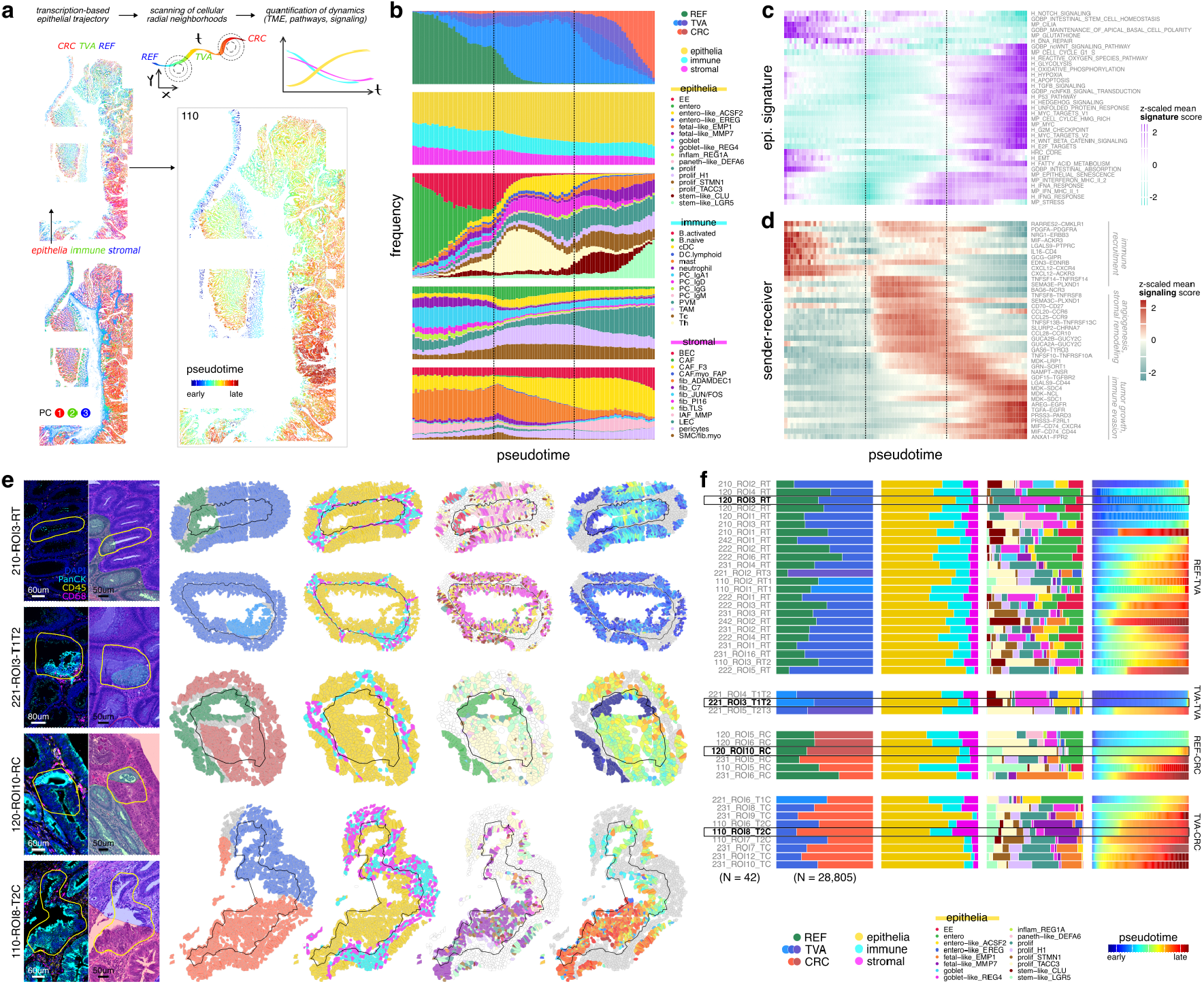
Epithelial trajectory and transition crypts. **(a)** Schematic illustrating transcription-based epithelial trajectory inferences and downstream analyses via the quantification of (physical) cellular neighborhoods through (pseudo)time *t*; included are spatial plots of section 110 with cells colored by an RGB representation of principal components (PCs) 1-3, and of epithelial cells colored by PCs and pseudotime, respectively (c.f. Fig. S6 and Fig. S13a). **(b)** Frequency of histopathological regions (epithelia only), compartment-level and -wise subpopulations over pseudotime; for non-epithelia, values correspond to the proportion of cells within a 50um radial neighborhood (c.f. Fig. S13b). **(c)** Epithelial gene set signature scores over pseudotime; y-axis ordered by hierarchical clustering. **(d)** Cell-cell communication over pseudotime; top-40 interactions according to (unscaled) value range. For (b-d), x-axes correspond to binned pseudotime and y-axis values are averaged by bin (see Methods). **(e)** Exemplary transition crypts between histopathological regions, namely, (fttb) REF and TVA, different TVAs, REF and CRC, TVA and CRC. Shown are (fltr) IF composite images and H&E stains, as well as spatial plots colored by histopathological region, compartment, epithelial subpopulation, and pseudotime (c.f. Fig. S14, Fig. S15 and Fig. S16). **(f)** Frequency of (fltr) histopathological regions, compartments, epithelial subpopulations, and pseudotime estimates across transition crypts; y-axis grouped by transition type, and ordered by average pseudotime.

Quantifying cellular state composition across pseudotime (Fig. 3b and Fig. S13b) demarcated three phases during which immune and stromal cells decrease in favor of epithelia. Early pseudotime was dominated by normal epithelial clusters (EE and enterocytes), PC IgA1, and fib ADAMDEC1. During mid-pseudotime, entero-, goblet-like and prolif epithelia emerged alongside IAFs, CAFs and TAMs. Stemand/or fetal-like populations dominated in the late-pseudotime phase. These two terminal populations exhibited mutually exclusive spatial distributions^[48]^ where one predominated over the other at different time points, in different donors, and in accordance with malignancy scores (c.f. Fig. 1c). Along the same axis, we observed higher proportions of Tc in less malignant samples, and higher proportions of TAMs in the most malignant ones.

We additionally observed functional and metabolic pathways shifts in epithelia that corresponded to different stages of carcinogenesis (Fig. 3c). In the early phase, cells showed enrichment of normal intestinal function and homeostasis pathways (cilia, maintenance of apical basal cell polarity, intestinal absorption, intestinal stem cell homeostasis, glutathione). As pseudotime progressed, we detected an enrichment of pathways known to drive cancer proliferation and progression (non-canonical and beta-catenin WNT, non-canonical NFKB, hedgehog, MYC, etc.), cell cycle and hypermetabolism (cell cycle G1-S, oxidative phosphorylation, glycolysis, G2M checkpoint, E2F, fatty acid metabolism), cell damage and stress (ROS, hypoxia, apoptosis, p53, senescence, unfolded protein response, stress), and increased expression of immune-related signatures such as IFN signaling. Late pseudotime phases showed an enrichment of signatures related to malignancy and invasion/metastasis: HRC core and EMT.

Cell-cell communication analysis reflected spatially-resolved signaling shifts among all compartments as the lesion progressed (Fig. 3d). Highly expressed interactions during early pseudotime (RARRES2–CMKLR1, IL16–CD4, CXCL12–CXCR4/ACKR3) suggested immune recruitment together with the modulation of inflammation (MIF–ACKR3, LGALS9–PTPRC). The upregulation of PDGFA/PDGFD–PDGFRA/B and SEMA3E/3C–PLXND1 during mid-pseudotime pointed to an engagement of fibroblasts, pericytes and endothelial cells to promote ECM remodeling and vascularization, while signaling via EGR ligands (NRG1–ERBB3, EREG/AREG–EGFR, TGFA–EGFR) indicated an early epithelial proliferation. The late pseudotime phase was dominated by growth factor and oncogenic signaling, with a strong increase in the EGFR ligands AREG, EREG, TGFA, as well as GDF15–TGFBR2, MDK–SDC1/SDC4 and NAMPT–INSR. In addition, LGALS9–CD44, MIF–CD74/CD44, CD70–CD27, TNFSF13/13B/10–TNFRSF13C/F10A and ANXA1/FPR2 indicate immune suppression and immune escape mechanisms.

Most intriguingly, the transitions from REF mucosa to adenoma and carcinoma described at the tissue-wide scale, defining the overarching tissue architecture, could also be mapped at the level of individual crypts. Epithelial crypt fusion (two neighboring crypts fusing into a single daughter crypt) has been described as a homeostatic mechanism in the human colon^[49]^, and has also been suggested to play a role in premalignant proliferative lesions^[50]^ from a genetic perspective. Here, we introduce the term ‘transition crypts’ to describe changes in cell morphology within the same intestinal crypt unit – a potential result of previous fusion or abrupt transformation events. 42 transition crypts were manually selected and annotated (Fig. S14, Fig. S15 and Fig. S16). Specifically, we identified 24 transitions from REF to TVA (RT), 6 from REF to CRC (RC), 9 from TVA to CRC (TC), and 3 transitions between differentially malignant TVAs (TT); see Fig. 3e,f. Single-crypt transitions comprised as few as ∼60-2,000 cells (600 on average). Crypts that transitioned away from REF (RT and RC) showed an abrupt drop in PanCK staining, while REF-free crypts (TT and TC) had overall dimmer signal (c.f., IF images in Fig. 3e and Fig. S14).

To further characterize these transition events, we quantified the epithelial, immune and stromal composition of each individual crypt within a 50um distance from the crypt boundary; see Fig. 3e and Methods. Almost all RC and RT crypts included REF clusters (EE, enterocytes, and goblet cells). In contrast, crypts transitioning to CRC (RC and TC) included stem-like, proliferative, and fetal-like epithelial clusters (Fig. 3e,f). In line with our previous observations at the tissue level, the immune fraction tended to decrease in transition crypts that lacked reference mucosa.

Next, we incorporated trajectory inference analysis to delineate individual transition crypt dynamics (right-most columns in Fig. 3e,f). Trajectories that were inferred at a tissue-wide scale, effectively recapitulated histologically-defined transitions at the single-crypt level. Here, crypts either underwent abrupt transitions, marked by sudden changes in both cluster composition and pseudotime assignment (e.g., 120-ROI10-RC), or exhibited more gradual transitions (e.g., 210-ROI3-RT). In general, RT crypts demonstrated greater epithelial heterogeneity compared to other transitions, suggesting distinct modes of tumor evolution that range from progressive changes to abrupt shifts in cellular identity. Abrupt transitions from reference mucosa to CRC (RC) tended to initiate from crypts that were already advanced in their pseudotime trajectories, suggesting a transition from precursor hyperplastic states. In line with this, carcinoma cells within these transition crypts were not fully advanced in their malignant transformation trajectory, pointing to a rapid yet incomplete malignant conversion.

In conclusion, we describe transition crypts as single functional units that show neoplastic transformation evidenced by cell morphology, cellular composition, transcriptional identity, and supportive niches.

### Tracking primary growth to spread from vascular invasion to metastasis

Detailed histopathology inspection further identified nine events of lymphovascular invasions (LIs) within CRC domains across several donors (120, 221, 232 and 242; Fig. S19 and Fig. S20). LIs were observed as clusters of epithelial cells located within the lumen of blood or lymphatic vessels. Invasive epithelial populations were predominantly composed of fetal-like MMP7 clusters (Fig. 4a,b), aligning with previous reports suggesting that MMP function plays a key role in cellular invasion, migration, and EMT^[51]^. To characterize such invasion events at a niche level, we also annotated blood vessels (BVs) located within CRC regions, but lacking intraluminal tumor cells (3 per section; Fig. S17 and Fig. S18); respective examples are shown in Fig. 4a. While the immune and stromal composition around control BVs was highly heterogeneous, the perivascular niche surrounding LIs was largely dominated by TAMs and CAF.myo FAP. In line with this, the supportive function of these specific immune and stromal cell states has been previously associated with lymphovascular invasion, poor histological differentiation, and lymph node metastasis ^[52,53]^, here further supported at the spatially-resolved, single-cell level.

**Figure 4.**
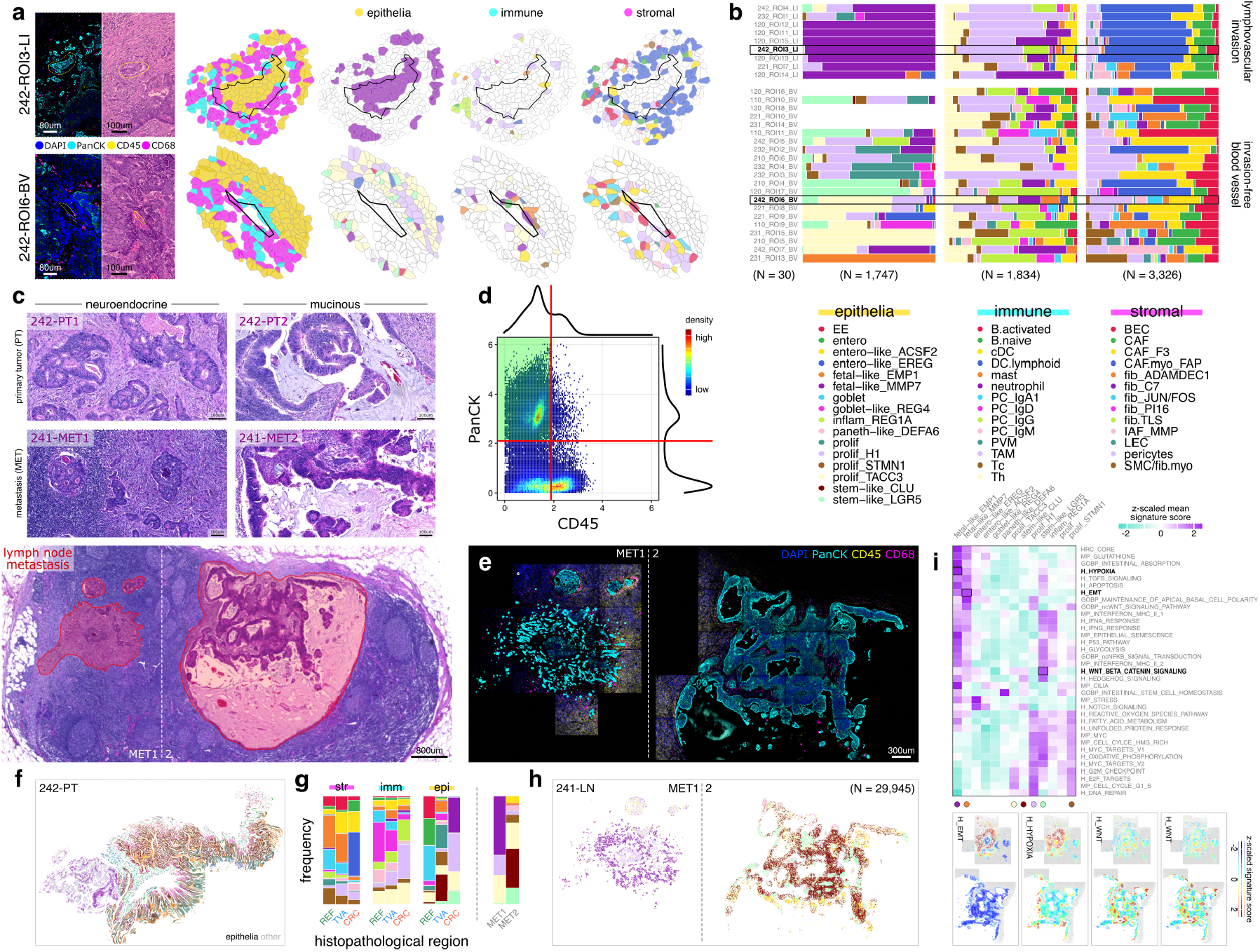
Lymphovascular invasions and lymph node metastases. **(a)** Selected lymphovascular invasions (LI; top) and invasion-free blood vessels (BV; bottom). Shown are (fltr) immunofluorescence (IF) composite images and H&E stains, as well as spatial plots colored by compartment-level and -wise subpopulation assignment (c.f. Fig. S17, Fig. S18, Fig. S19 and Fig. S20). **(b)** Frequency of compartment-wise subpopulations across LIs (top) and invasion-free BVs (bottom); y-axis ordered by hierarchical clustering of epithelia. **(c)** H&E stain overlaid with histopathological annotation of lymph node metastatic nodules. Higher magnification of both metastases (section 241) and comparison with primary CRC regions from the same donor (section 242), showing distinct neuroendocrine or mucinous growth patterns (c.f. Fig. S2). **(d)** Scatter plot of PanCK and CD45, including density-based threshold estimates; axis values correspond to mean per-cell intensities, arcsinh-transformed with a cofactor of 200. **(e)** IF composite image (section 241; lymph node metastasis). **(f)** Spatial plots with cells colored by epithelial subpopulation (section 242; primary tumor). **(g)** Compartment-wise subpopulation frequencies across section 242’s histopathological regions, and of epithelial subpopulations across section 241’s metastatic nodules. **(h)** Spatial plot with cells colored by epithelial subpopulation (section 241). **(i)** Gene set signature scores across epithelial subpopulations (data are across all colon samples), alongside spatial plots with cells colored by selected gene set signature scores (section 241).

One donor with LIs in the primary tumor (242-PT, the most malignant case in our study; c.f. Fig. 1c and Table S1) also presented with metastases (METs) in a draining lymph node (241-LN). Whole-slide H&E stainings of the PT and METs identified notable tumor heterogeneity, with two distinct tumor growth patterns present at both sites: solid, undifferentiated tumor cells with neuroendocrine differentiation (242-PT1 and 241-MET1), and a more differentiated, mucinous region (242-PT2 and 241-MET2); see Fig. 4c. The PT’s mucinous component (242-PT2) was not captured by the CosMx instrument (outside the scanning area defined *a priori*), limiting transcriptional characterization of the tumor cells in that region (Fig. S2).

We next used the mean fluorescence intensities of PanCK and CD45 stainings from the LN section to subset tumor cells from the two morphologically distinct metastatic nodules (CD45-PanCK+; see Fig. 4d,e and Methods). Using supervised clustering with *InSituType*^[31]^ anchored to the primary tumor profiles (see Methods), we found that MET1 and MET2 exhibited markedly distinct transcriptional programs^[54]^. On the one hand, MET1 was enriched in fetal-like MMP7 and prolif H1, mirroring the cellular composition of the invasive CRC domain in the primary site (232-PT, Fig. 4f-h) and the tumor cells present in LIs (Fig. 4a,b). On the other hand, MET2 was composed primarily of stem-like LGR5 and CLU populations, also observed at the tumor core in the primary tumor. Correspondingly, scoring relevant gene signatures across transformed epithelial cells from the PT and LN (Fig. 4i) detected elevated hypoxia and EMT in MET1, previously assigned to fetal-like states. By contrast, MET2 showed increased WNT signaling (promoted by LGR5), pathways activated in stem-like populations.

These results suggest that the histopathologically and transcriptionally distinct metastases in the draining LN arose simultaneously or subsequently from different CRC subpopulations of the primary tumor. Alternatively, the same tumor origin gave rise to divergent tumor states at proximal metastatic sites^[55,56]^.

## Discussion

We present a single cell- and spatially-resolved architectural map of 3.5M cells in colorectal tissues that recapitulate the malignant transformation of normal mucosa to adenoma and carcinoma stage, combining transcriptome-wide spatial transcriptomics (WTx CosMx SMI) with single-nucleus sequencing (snPATHO-seq) and digital histopathology.

Considering these orthogonal data views proved valuable in resolving ambiguous regions for the pathologists, and offered molecular confirmation of malignant features that were not unequivocally identifiable through histology. Our bottom-up and top-down analysis strategies, anchored in trajectory inference, leveraged spatial tissue structure at all scales: from single cells, via local interactions and functional units (cellular niches, transition crypts, vessel structures), to tissue-wide scale. Together, this allowed for a wholesome understanding of the molecular and cellular dynamics underlying CRC emergence and progression, not only considering the neoplastic transition of epithelial states, but also the microenvironmental and architectural traits that support tumor growth and dissemination.

Applying latest transcriptome-wide gene panels (WTx) let us incorporate spatial information with analytical avenues previously hampered by limited plexity of imaging-based ST data; for instance, trajectory inference^[45,57]^ and the systematic interrogation of annotated gene sets^[58,59]^ and ligand-receptor interactions^[60]^. Further, our data-driven approaches were guided by the integration of histopathology annotations, particularly valuable in sections where normal crypts, adenomas and invasive carcinomas coexisted.

We identified 43 transcriptionally and spatially distinct subpopulations spanning epithelial, immune, and stromal compartments. Trajectory analysis computed on 16 epithelial states accurately recapitulate the malignant CRC progression in all samples. The cellular interactions viewed along epithelial pseudotime captured the premalignant transitions to adenoma (e.g. epithelial proliferation, immune recruitment and ECM remodeling) and subsequent evolution into carcinoma (e.g. tumor growth and immune evasion). Notably, the shift from TVA to CRC occurred as a fate determination, which led to higher epithelial homogeneity in favor of two key tumor populations: i) a fetal-like MMP7 state, marking invasive fronts and enriched in EMT, TGFB and CAF pathways/metaprograms; and, ii) a stem-cell LGR5 population enriched in WNT signaling and confined to the tumor core.

While stem-like LGR5 tumor cells were densely packed and engaged in homophylic interactions, suggestive of niche/core building, the fetal-like MMP7 tumor cell state was found in looser and heterogeneous microenvironments with an immunosuppressive and pro-invasive molecular and cellular composition (enriched in CAF.myo FAP and TAMs; reduced Tc cells). Previous reports have linked CAF.myo FAP, SPP1+ TAMs and low Tc infiltration to poor prognosis and therapy response rates in CRC^[61]^. In our study, these organizational shifts also aligned with histological malignancy grading, indicating that transcriptional reprogramming and niche reshaping are tightly linked to tumor progression.

Further tracking tumor progression identified small clusters of fetal-like MMP7 tumor cells located inside vessels and surrounded by CAF.myo FAP. One donor had two draining lymph node metastases with distinct growth patterns and transcriptional profiles, one of them resembling the corresponding primary tumor. Spatially-separated metastases with divergent epithelial subtypes (at both secondary and primary sites) suggest seeding events that differ either in their origin or timing, and with potentially different metastatic behavior and therapeutic sensitivity.

Adenomas were the most heterogeneous stage. The presence of a CLU+ revival stem cell population in adenomas suggests the activation of regenerative programs prior to malignant transformation. Further, the identification of an EMP1+ epithelial state (epi.fetal-like EMP1) in the TVA domain, closely resembling HR EMP1+ cells described as the origin of metastatic recurrence by Cañellas-Socias *et al*.^[48]^, is indicative of clinically-relevant populations related to patient relapse being present already at premalignant stages of CRC development.

We acknowledge the limitations of this study, in particular the bias of analyzing tumors in two dimensions (2D) and using gene expression as the major readout for cell phenotyping. Viewing tumors as corporal structures has already been shown to derive new insights into CRC biology^[26]^, by resolving low-plex panels directly in 3D^[62]^, or via the virtual reconstruction of tissue blocks^[63]^. In addition, multi-modal spatial omics are becoming available ^[64,65]^, laying the ground stones for deciphering also regulatory dependencies of multi-cellular systems, such as tumors. Meanwhile, the application of machine learning and artificial intelligence in biomedical research is gaining traction and will be key in spatial biology, data analysis and interpretation^[66]^. Thus, both methodological and biotechnological leaps promise to shed new light on CRC and research at large in the close future.

## Data availability

All data have been deposited on Zenodo at the following DOIs. H&E stains are available at 10.5281/zenodo.15548055. Flat files of the CosMx SMI data (FOV placement, count matrices, cell metadata, segmentation boundaries) are available at 10.5281/zenodo.15550908. Napari inputs (Zarr stores of IF markers, FOV and segmentation labels, single-molecule RNA targets), including histopathological domain and ROI annotations, are available at 10.5281/zenodo.15556499 (11), 10.5281/zenodo.15556609 (12), 10.5281/zenodo.15556207 (21), 10.5281/zenodo.15555405 (22), 10.5281/zenodo.15552301 (23), and 10.5281/zenodo.15551924 (24); one file batch per slide (due to file size limitations, some files for sections 11 and 12 are placed at 10.5281/zenodo.15585754).

We also make available *SingleCellExperiment* (.rds) and *AnnData* objects (.h5ad; written using *zellkonverter* ^[67]^), as well as language-agnostic *alabaster* file artifacts (written using *alabaster*.*sce*^[68]^) that, for each section, gather processing and analysis results for the CosMx SMI data; these are available at 10.5281/zenodo.15574384. Also included are (un)filtered *cellranger* outputs for the snPATHO-seq data, as well as the lowand high-resolution reference profiles derived thereof, which were used for clustering of the CosMx SMI data.

## Code availability

All analyses were run in R v4.4.2^[69]^ with Bioconductor v3.20^[70]^. The computational workflow was implemented using *Snakemake* v8.20.5^[71]^, with Python v3.12.6. Data were handled using the *SingleCellExperiment* container^[72]^ with a delayed backend (using *HD5Array* ^[73]^). Visualizations were generated using custom *ggplot2*^[74]^ code (R plots), or via export from QuPath^[75]^ (H&E stains) and Napari^[76]^ (IF images). *Snakemake* workflow and underlying R code are accessible at https://github.com/HelenaLC/WTx-TVA.

## Acknowledgement

H.L.C. is funded by the Swiss National Science Foundation (project grant number 222136). A.P.R. receives funding from the MCIN/AEI/10.13039/501100011033 and FSE+ (RYC2022-035848-I) and the MICIU/AEI/10.13039/501100011033/ FEDER/UE (PID2023-148687OB-I00). H.H. receives funding from the European Union’s H2020 research and innovation program (848028), from the European Research Council (ERC) (810287), from the Ministerio de Ciencia e Innovació n (MCI) (PID2020-115439GB-I00 and PLEC2021-007654), from the LaCaixa Foundation (HR22-0031 and HR22-0172), from the Generalitat de Catalunya through the Suport Grups de Recerca AGAUR (2021-SGR), from ERA-NET Neuron/Ministerio de Ciencia e Innovació n (MCI) (PCI2022-133012), and from an ASPIRE Award from The Mark Foundation for Cancer Research and the Scientific Foundation of the Spanish Association Against Cancer.

## Authors’ contributions

H.L.C. performed computational analyses, generated figures, and wrote computational methods. I.R. performed histopathological classifications, generated IF- and H&E-based images, and wrote corresponding methods. A.P.R., I.R. and H.L.C. interpreted the data and results, conceptualized and drafted the manuscript, with contributions from all authors. Z.H., Y.H., H.P. and S.T. selected and prepared sections, and aided in the interpretation of results. G.C. performed snPATHO-seq experiments. J.B. conceived and implemented the chemistry expansion to whole transcriptome levels. A.H., R.L., K.Y., M.W., M.V., D.M. and M.L.H. worked to coordinate, execute, and process the CosMx WTx experiments. E.M. aided in the spatial analysis with Napari. H.H. acquired funding, helped draft the manuscript, and conceptualized the study together with S.T. All authors read and approved the final manuscript.

## Competing interests

H.H. is co-founder and Chief Scientific Officer of Omniscope, a Scientific Advisory Board member at NanoString/Bruker and Mirxes, a consultant for Moderna and Singularity, and has received honoraria from Genentech. A.H., R.L., M.W., M.V., K.Y., D.M., E.M., M.L.H. and J.M.B. are employees of Bruker.

## Methods

### Sample collection and details

This study was approved by the Institutional Review Board (study number S66460), and all patients provided written informed consent. Seven treatment-naïve CRC patients were enrolled. Eight primary colon tissue sections and one lymph node section were collected intraoperatively, immediately placed on ice, and transported to the pathology laboratory within 20 minutes. Specimens were fixed in 10% neutral buffered formalin for 24 hours, embedded in paraffin, and cut into 5 um sections. Consecutive sections were obtained, and two 25 um sections from each block were reserved for single-nucleus RNA sequencing. Adjacent 5 um sections were stained with hematoxylin and eosin (H&E), while parallel 5 um sections were used for CosMx spatial transcriptomics. Both H&E and CosMx slides were shipped to Seattle under room temperature. See Table S4 for clinical and genetic features of the specimens.

### Histopathology classification

#### Colon annotation: REF, TVA and CRC regions

Tissue sections stained with H&E were first annotated manually in a blinded-manner using QuPath^[75]^ (v0.5.1). Two pathologists independently annotated H&E stained sections under a double-blind protocol. For each donor, consecutive immunofluorescence digital images obtained by the CosMx instrument from sections stained with nuclear (DAPI) and cell segmentation markers (CD45, CD68, PanCK; c.f. Fig. S1) were annotated in Napari ^[76]^, using previous H&E annotations as guidance. Region-wise classifications, corresponding criteria, and additionally observed characteristics are detailed in Table S3 and Fig. S2.

Colon mucosa was classified as reference (REF), premalignant polypoid tubulovillous adenoma (TVA) or malignant colorectal cancer (CRC). If distinct areas of TVA or CRC were detected (sections 110, 120 and 221), separate annotations were made in order of malignancy; e.g., CRC2 is more malignant than CRC1.

The healthy or reference-like mucosa (REF) showed a preserved normal crypt architecture, and no tumor epithelial proliferation. The lamina propria was expanded with minimal to high infiltration of inflammatory cells. In all samples, REF regions showed an increase in the number of cells with normal morphology (hyperplasia), specially those close to TVA or CRC growth.

TVA and CRC classification criteria and annotations were performed according to the 2019 World Health Organization (WHO) guidelines^[2]^. Conventional premalignant adenomas are benign polypoid lesions characterized by a dysplastic epithelium. Histologically, they can be differentiated into three different subtypes (tubular, tubulovillous, and villous)^[77]^ depending on their growth architecture. Here, all adenomas were consistent with a mixed pattern (tubulovillous adenoma, TVA), with a villous component between 25-75%, although 110 TVA3, 210 TVA and 242 TVA regions presented a more evident villous component. TVAs were then classified as lowor high-grade dysplasia. The vast majority of sections showed low-grade dysplasia, except for two regions (110 TVA3 and 242 TVA) that showed features consistent with high-grade dysplasia (irregular crowded glands, cribriform architecture, or intraluminal necrosis).

#### CRC classification criteria

Colorectal carcinoma (CRC) correspond with a malignant proliferation of epithelial cells growing from the mucosa and effacing deeper layers. Different CRC subtypes recognized by the WHO can be determined at the histopathological level^[78]^. The most common, colon adenomatous adenocarcinoma, is characterized by glandular formation, often cribriform and filled with necrotic debris. On the other hand, mucinous adenocarcinoma is a special subtype of CRC defined by more than 50% of the tumor composed of extracellular mucin with floating epithelial glands, occasionally with undifferentiated morphology ^[79]^. The CRCs of sections 120 and 221 were classified as adenomatous adenocarcinoma, and CRCs from sections 210, 231 and 232 were classified as mucinous adenocarcinomas. Additionally, section 110’s CRC region showed proliferation of malignant colorectal cancer cells growing in a villous-like pattern resembling small intestine; this region was designed as villous CRC. A fourth subtype of CRC was identify in sample 242, which presented morphologic features suggestive of an adenomatous CRC with neuroendocrine differentiation (NED), described by WHO guidelines as an aggressive and undifferentiated neoplasm, and composed of solid trabecular nests of malignant cells with large cytoplasm, prominent nucleoli, frequent mitosis, and abundant necrosis^[78]^.

In addition to histological subtypes, CRC regions were graded according to their glandular cell differentiation and depth of tumor invasion. Regarding differentiation, the WHO classifies CRC as: well-differentiated (G1) when showing more than 95% of gland formation; moderately-differentiated (G2) when gland formation ranges between 50-95%; and poorly-differentiated (G3) when gland formation is lower than 50%. In this study, 3/8 CRC were graded as G1, 1/8 as G2, and 3/8 as G3. At a finer level of detail, isolated clusters of malignant undifferentiated tumor cells were observed in CRCs of sections 210 and 242.

CRC classification based on tumor invasion follows the American Joint Committee on Cancer’s tumor, node, metastasis (TNM) system, where the T category describes the depth of tumor infiltration^[1]^: remains confined to the muscularis mucosa (T0); further invades the muscularis propria (T1); extends into the submucosa (T2); penetrates through the muscularis propria into the outermost layers of the colon (T3); or, grows through the entire wall of the colon (T4). In this study, 1/8 CRC were staged at T0, 1/8 at T1, 5/8 at T2, and 1/8 at T3. Tissue orientation in 221 CRC, marked with an asterisk in Table S3, impaired an accurate invasion scoring.

#### CRC-associated features

Further cancer-associated histopathological features, such as desmoplastic reaction (DR), angiogenesis, necrosis or tertiary lymphoid structures (TLS), are also summarized in Table S3. DR is characterized by the dense proliferation of spindle-shaped stromal cells surrounding the tumor, indicating active remodeling of the TME^[80]^. All CRC tissue samples staged at T2 or higher presented evident DR. Angiogenesis, development of new blood vessels from precursor endothelial cells, is occurs during cancer development and recognized as one of the principal features of CRC ^[81]^. In the present report, four CRC regions showed this feature (120, 221, 232 and 242) which directly correlated with higher differentiation, invasion and inflammation. Another histological finding evaluated was the presence of TLS – ectopic clusters of immune cells found within the TME correlating with favorable clinical outcomes ^[82]^; four donors showed evident TLS formation in CRC regions.

In addition, certain areas within the CRC domain exhibited focal or multifocal disruptions of normal architecture and replacement with dense fibrin deposits, hemorrhagic components, and accumulations of cellular debris, all indicative of necrosis. Notably, these necrotic zones were often situated in the apical portion of the tumor mass, which may be attributable to rapid tumor expansion.

Immune infiltrates were thoroughly assessed in all regions (REF, TVA, and CRC) from each sample. The severity of inflammation was graded using a 5-point scoring system ranging from none (0) to minimal (1), mild (2), moderate (3), and high (4) based on the extent of immune cell infiltration surrounding epithelial cells. Additionally, the predominant immune cell types were identified and classified, distinguishing between mononuclear (MN) and polymorphonuclear (PMN) cells, and thus providing a comprehensive analysis of the immune landscape in each case.

### Regions of interest (ROIs)

#### Transition crypts

Besides REF, TVA and CRC tissue domains, some biologically relevant regions of interest (ROIs) were also described and annotated in H&E stained tissue sections (using QuPath), and used to guide annotations of IF stained CosMx slides (using Napari). Single intestinal crypts in which the epithelial compartment transitioned between distinct domains, which exhibited well-oriented tissue architecture with clear basement membrane continuity, were annotated. Within these crypts, notable transitions in epithelial cellular morphology were observed, including abrupt changes from REF to TVA, REF to CRC, and TVA to CRC. Although 46 individual crypts were first annotated in H&E stains, four of these did not preserve clear basement membrane continuity in consecutive IF slides, so that they were excluded from downstream analysis. In sum, we identified 24 transitions from REF to TVA (RT), 6 from REF to CRC (RC), 9 from TVA to CRC (TC), and 3 transition between different TVA regions (TT).

#### Lymphovascular invasions

Certain CRC regions exhibited a limited presence of undifferentiated tumor cells within the lumen of blood or lymphatic vessels; this phenomenon is called lymphovascular invasion (LI). Nine LIs from five different donors were annotated. Control blood vessels (BVs) located within the CRC regions but without any evidence of tumor invasion were also annotated for comparison (three from each CRC region, 21 in total).

#### Metastatic lymph node section

One lymph node (LN) sample was provided in the cohort, originating from the same donor as section 242. LN was annotated following the same workflow as described for colon tissue sections (using QuPath and Napari). Various metastatic nodules were identified, composed of heterogeneous proliferation of large pleomorphic tumor epithelial cells that replace and distort the normal lymphoid architecture. Two distinct growth patterns were observed: a solid trabecular pattern with numerous undifferentiated clusters and surrounded by abundant desmoplastic reaction (referred to as MET1), and an organized and well-differentiated tumor forming glandular and tubular structures, frequently dilated with abundant mucus admixed with cellular debris and fibrin (named MET2).

### Single-nucleus RNA sequencing (snPATHO-seq)

#### Sample preparation and data acquisition

Nuclei suspension preparation was performed following the snPATHO-Seq v7 protocol from Vallejo *et al*.^[83]^. Briefly, two 25 um-thick formaldehyde-fixed, paraffin-embedded (FFPE) tissue sections were washed three times with 1 mL xylene for 10 minutes each to remove paraffin. Tissue rehydration was carried out using sequential ethanol washes: 2 × 1 mL of 100%, followed by 1 × 70%, 1 × 50%, and 1 × 30% ethanol (1 minute each), and finally 1 × 1 mL RPMI-1640 medium (Gibco). Tissues were digested in 1 mL of RPMI-1640 supplemented with mg/mL Liberase TH (Sigma-Aldrich) for 60 minutes at 37 °C in a thermomixer set to 800 rpm. Post-incubation, samples were treated with Ez Lysis Buffer (Sigma-Aldrich) to release nuclei from partially digested tissues. Homogenization was performed using a plastic pestle in 250 uL Ez Lysis Buffer supplemented with 2% BSA (Miltenyi Biotec) and 1 U/uL RNase Inhibitor (Roche). An additional 750 uL of the same buffer was added, and samples were incubated on ice for 10 minutes. The suspension was filtered through a 70 um filter (PluriStrainer) and washed once with 800 uL Ez Lysis Buffer + 2% BSA + 1 U/uL RNase Inhibitor, followed by two washes with 0.5× PBS (Thermo Fisher Scientific) + 0.02% BSA. All centrifugation steps were performed at 850 × RCF for 5 minutes at 4 °C. After washing, nuclei were resuspended in 500 uL of 0.5× PBS + 0.02% BSA, filtered through a 40 um filter (PluriStrainer), and counted using the Luna-FL cell counter (Logos Biosystems) after AO/PI staining.

snRNA sequencing was conducted using the 10X Genomics Chromium Flex platform (NextGEM version). For each sample, 150,000–650,000 nuclei were used as input for hybridization with the whole human transcriptome probe kit (10X Genomics, PN-1000475), performed at 42 °C for 22 hours in a 4-plex configuration, following the manufacturer’s user guide (CG000527 Rev D). After hybridization, samples were washed individually, resuspended in 500 uL buffer, filtered through a 30 um filter (Sysmex CellTrics), and recounted using the Luna-FL system. Samples were pooled in equimolar ratios, and the final pool was re-quantified before loading onto the Chromium X instrument (10X Genomics), targeting recovery of 40,000 nuclei. Library preparation followed the manufacturer’s instructions, using 12 cycles for indexing PCR. The quality and quantity of final libraries were assessed using the Agilent Bioanalyzer system (Agilent Technologies). Sequencing was performed on the Illumina NovaSeq platform, targeting ∼20,000 reads per nucleus with the following configuration: 28 bp (Read 1) + 10 bp (i7 Index) + 10 bp (i5 Index) + 90 bp (Read 2).

### Computational data processing and analysis

#### Decontamination

Following log-library size normalization (using *scater* ^[84]^), we first selected for features that were among the top-2,000 most highly variable (using *scran*^[85]^; blocking by patient) in more than one sample (N=2,797). These were used for principal component analysis (PCA) using *irlba*^[86]^. The first 30 PCs were then corrected for batch effects between runs using *harmony* ^[87]^.

Next, a shared-nearest neighbor (sNN) graph was computed with *distance=“jaccard”* and *k=30*, and clustered using the Louvain community detection algorithm^[88]^ with *resolution=1*, yielding 20 clusters. These preliminary cluster assignments were passed to *decontX* ^[89]^, alongside raw data on the (unfiltered) barcodes, in order to correct for ambient RNA contamination.

#### Quality control

For each sample, we deemed as outliers cells below 1.5 MADs in total counts or uniquely detected features, and above 3 MADs in the percentage of mitochondrial counts; besides, only genes with more than one count in at lest 20 cells were kept. Taken together, 14,366 genes and 115,869 cells were retained for downstream analysis.

Data were then reprocessed analogous to above (i.e., log-normalization of decontaminated counts, feature selection, batch correction, and sNN graph construction), and clustered using the Leiden community detection algorithm^[90]^ with *resolution=1*, optimizing for modularity. Cluster assignments were passed to *scDblFinder* ^[91]^, grouping by patient; cells classified as doublets, as well as one cluster with relatively high doublet scores after doublet removal, were left unassigned.

#### Annotation

Clusters were annotated into 16 broad subpopulations based on results from differential expression (DE) analysis (using *scran*^[85]^); namely, IgA/G/M plasma, B, T, mast, endothelial and smooth muscle cells, as well as two myeloid, two fibroblast, and four epithelial subpopulations.

Based on level 1 annotations, cells were split into epithelial (four subpopulations), immune (IgA/G/M plasma, B, T, mast cells, and two myeloid subpopulations), and stromal (endothelial and smooth muscle cells, and two fibroblast subpopulations) compartments. For each compartment, data were processed analogous to above (feature selection, batch correction), except: the sNN graph was built using *k=20*, and Leiden clustering was performed with a *resolution* of 1.8, 1.4 and 1.0 for the epithelial, immune and stromal compartment, yielding 18, 18 and 20 clusters, respectively; these were annotated into 16, 14 and 13 subpopulations, respectively.

#### Reference profiles

For level 1 annotations, genes ranked among the top-200 DE genes in any comparison (N=4,049) were selected. For level 2 annotations, DE genes were extracted separately for each compartment. Only genes that ranked among the top-200 genes in at least 2 comparisons (N=3,857 for epithelia, 1,872 for immune, and 1,534 for stromal cells) were considered.

Lastly, pseudobulks were computed by summing counts per sample and cluster, normalizing for library size, and averaging normalized counts across samples (using *scater* ‘s *aggregateAcrossCells()* function). The respective feature subsets were used as reference profiles for *InSituType* clustering of the CosMx SMI data, as well as PCA, etc.

### CosMx Spatial Molecular Imager (SMI)

#### Sample preparation and data acquisition

To perform *in situ* hybridization on five-micron tissue sections, slides were baked in an oven at 60 °C overnight. Tissue sections were dewaxed twice in xylene (Millipore) for 5 min, twice in ethanol (Pharmco) for 2 min and then the slides were baked at 60 °C for 5 min. Tissue sections were subjected to the target retrieval step using CosMx Target Retrieval Solution and heated at 100 °C in a pressure cooker for 15 min. After target retrieval, tissue sections were rinsed with diethyl pyrocarbonate (DEPC)-treated water (DEPC H2O, ThermoFisher), washed in ethanol for 3 min and dried at room temperature for 30 min. On the dried slide, an adhesive incubation frame was placed around the tissue section. Tissue was then digested with Proteinase K (ThermoFisher) at 3 ug/ml concentration at 40 °C for 30 min. Tissue sections were rinsed twice with DEPC H2O, incubated in 1:100 diluted fiducials (Bangs Laboratories) in 2X saline sodium citrate and Tween (0.001% Tween 20, Teknova) for 5 min at room temperature and washed with 1X PBS (ThermoFisher) for 5 min. After digestion and fiducial placement, tissue was fixed with 10% neutral-buffered formalin for 1 min to maintain soft tissue morphology, washed twice with Tris-glycine buffer (0.1 M glycine (Sigma), 0.1 M Tris-base (FisherScientific), in DEPC H2O) for 5 min and then washed with 1X PBS for 5 min. Fixed tissue was blocked using 100 mM Nsuccinimidyl (acetylthio) acetate (NHS-acetate, ThermoFisher) diluted in NHS-acetate buffer (0.1 M NaP + 0.1% Tween pH 8.0 in DEPC H2O) for 15 min at room temperature and washed in 2X saline sodium citrate (SSC) for 5 min.

The (pre-commercial) CosMx Human WTx panel consisted of designed 37,890 *in situ* hybridization (ISH) probes targeting 19,867 human genes covering *>*99.5% of the annotated protein-coding genes curated by the HUGO Gene Nomenclature Committee (HGNC15) for which there was available sequence in NCBI RefSeq16, as described in Khafizov *et al*.^[16]^.

CosMx WTx ISH probes were prepared by incubation at 95 °C for 2 min and immediately transferred to ice. The ISH probe mix (1 nM ISH probe, 40% formamide, 2.5% dextran sulfate, 0.2% BSA, 100 ug ml–1 salmon sperm DNA, 2× SSC, 0.1 U ul–1 `SUPERase·In` (ThermoFisher) in DEPC H2O) was then pipetted into the frame seal, and an incubation frame cover was applied. Hybridization occurred at 37 °C overnight after sealing the chamber to prevent evaporation. Following ISH probe hybridization, the incubation frame cover was removed, and tissue sections were washed twice in a buffer comprising 50% formamide (VWR) in 2X SSC at 37 °C for 25 min, then washed twice with 2X SSC for 2 min each at room temperature. Next, slides were incubated with CosMx Nuclear Stain for 15 min at room temperature, then washed in 1X PBS for 5 min. Tissue was then incubated with CosMx Cell Segmentation Mix for 1 h at room temperature and washed three times in 1X PBS. The adhesive incubation frame was removed before applying a flow cell coverslip and loading the slides on the CosMx SMI instrument.

The *in situ* chemistry and imaging analyses were performed on a commercial-grade CosMx instrument, in two runs of two and four slides each (c.f. Table S1). Technical details on the imaging and microfluidics system can be found in Khafizov *et al*.^[16]^. Pre-bleaching was performed with configuration C (60” per FOV); cell segmentation utilized profile A, including nuclear diameter (7.2um), cell diameter (8.28um) and dilation (2.16um).

### Computational data processing and analysis

#### Quality control

For each section, we deemed as outliers cells below 3 median absolute deviations (MADs) in total RNA counts and counts per area (computed on the log-scale using *scater* ‘s *isOutlier()* function^[84]^), as well as cells above more than 10 means in negative probe and blank code counts. Furthermore, because lack of FOV stitching during cell segmentation can result in fractured and duplicated cells near FOV borders, we filtered out cells closer than 30px to any FOV border in order to mitigate potential artifacts in downstream analyses. Jointly, these criteria removed ∼5% of cells per section (c.f. Table S1). Henceforth, expression values denote log-transformed library size-normalized counts, computed with *scater* ‘s *logNormCounts()* function.

#### Label transfer

Reference profiles were computed by summing counts per sample and cluster, library size normalization, and then averaging normalized counts across samples (using *scater* ‘s *aggregateAcrossCells()* for aggregation and *logNormCounts()* for normalization with *log = FALSE*). Semi-supervised clustering was performed using *InSituType*^[31]^ following the authors’ recommendations; briefly: To account for platform effects, reference profiles were adjusted using *updateReferenceProfiles()* with per-cell background estimated as the fraction of total RNA counts stemming from negative probes. Cohorts (via *fastCohorting()*) were based on mean immunofluorescence stains (DAPI, CD45, PanCK and PanMem), as well as cell area and aspect ratio. Lastly, *insitutype()* was run with *n clusts* 6-12, yielding 11 *de novo* and 13 labelled clusters, which were annotated into 22 subpopulations.

#### Normal epithelial subclustering

Normal epithelial subpopulations (EE, entero and goblet) were subset and clustered as above, using reference profiles from Elmentaite *et al*.^[44]^. Specifically, *Pan-GI Cell Atlas* data were retrieved from https://www.gutcellatlas.org, and filtered to retain epithelia of the large intestine from control individuals only (∼60,000 of 1M cells). In addition, we simplified original annotations (9 subpopulations), merging two colonocyte and enteroendocrine subsets each, thus retaining 7 subpopulations for label transfer (BEST4+/enterocytes, enteroendocrine, goblet, stem, transit amplifying and tuft cells). Lastly, the top-100 differentially expressed genes were selected for each subpopulation (using *scran*’s *findMarkers()* function with *direction=‘up’*) and their union (1,296 features) was used to compute bulk reference profiles.

#### Cellular density and entropy

Compartment-wise densities were estimated by counting the number of epithelial/immune/stromal cells within a 50um radial neighborhood (using *RANN*’s *nn2()* function with *searchtype=“radius”*), z-scaling counts per section for visualization. Epithelial subpopulation densities (Fig. 2c) were computed in the same way, downsampling to at most 10,000 cells per section and subpopulation, such that each section contributes similarly. (Shannon) entropies (Fig. 2d-e) correspond to 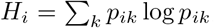, where *i* is a cell, *k* is a subpopulation, and *p*_*k*_ is the frequency of subpopulation *k* in cell *i*’s 50um radial neighborhood; for visualization, cells were downsampled as described above.

#### Signature scoring

Gene sets of interest were programmatically sourced from *MSigDB*^[58]^, and subset to features with matching gene symbols in our data. In total, we considered 47 epithelial, 26 immune and 18 stromal sets (81 unique) with 6-626 features (∼83 on average). Signature scores were computed with *AUCell*^[59]^, using *AUCell buildRankings()* (to build rankings) and *AUCell calcAUC()* with *aucMaxRank=400* (to calculate enrichment). In the resulting (sets × cells) matrix, values correspond to the fraction of genes in a given set that ranks among the top-400 genes in a given cell (i.e., values are proportions ∈ [0, 1]). These scores were averaged by cell subsets of interest for comparison across, e.g., histopathological regions, pseudotime bins, or subpopulations (c.f. Fig. 2f, Fig. 3c and Fig. 4i).

#### Niche analysis

To identify spatial context, we first quantified subpopulation frequencies among each cell’s local neighborhood; specifically, considering cells within a 50um radius (using *RANN*’s *nn2()* function with *searchtype=“radius”*). Notably, frequencies were ‘blinded’ to epithelial subpopulation identities, i.e., all epithelia were treated equally. The resulting (cells × clusters) matrix of proportions, pooled across sections, was then used as input to *k*-means clustering with *k=10*. We initially explored a range of clusterings (*k=8-15*); fewer clusters failed to recapitulate major histopathological domains, while more clusters tended to fragment niches into similarly composed ones. Thus, we chose 10 niches as a compromise between accurate resolution of tissue architecture, and ease of interpretability.

#### Cell-cell communication

Cell-cell communication analysis was performed using *COMMOT* ^[92]^, an optimal transport-based method that considers physical cell-to-cell distances and can account for competing sender/receiver signals as well as multimeter receptors. We considered 602 sender-receiver interactions (493 features, 121 pathways) available through *CellChatDB*^[60]^ for which all interaction partners were represented. The average cell area in our data was ∼83um^2^; here, we considered 50um as a threshold on signaling distance, which is about ten times an average (approximately circular) cell’s radius 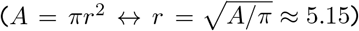.

#### Epithelial trajectory

We used *slingshot* ^[45]^ for spatially-unaware trajectory inference, using the same set of features underlying epithelial subclustering to perform PCA (using *scater* (with *BSPARAM = RandomParam()*). Specifically, the first 30 PCs and were passed to *slingshot()* (with *approx points = 100*) alongside histopathological region annotations as *clusterLabels*. Here, a given section’s REF region was used as start (*start*.*clus*), and the most malignant region as end point (*end*.*clus*). Pseudotimes used for downstream analyses and visualizations were computed with *averagePseudotime()*, then rescaling values between 0 and 1 using 1%- and 99%-quantiles as boundaries. In order to quantify pseudotemporal changes in microenvironment, cell-cell communication, and transcriptional signatures, etc., we grouped pseudotimes into ∼100 equally sized bins, and averaged values of interested (e.g., subpopulation frequencies) across a given epithelial cell’s 50um radial neighborhood.

#### Regions of interest (ROIs)

Transition crypts, invasion-free blood vessels (BVs), and lympho-vascular invasions (LIs) were first subset using Napari-based polygonal selections, spatially aligning coordinates to the CosMx SMI data, and subsetting cells whose centroids fall within a given ROI (using *point*.*in*.*polygon()* of the *sp* package^[93]^). Next, a concave hull was computed (using *concaveman*^[94]^), followed by a 50um polygonal expansion (using *st buffer()* of the *sf* package^[95]^). Buffered polygon coordinates were then used to identify cells whose centroids fall within a given ROI; these were used for visualization and quantification (c.f. Fig. 3e,f and Fig. 4a,b).

#### Lymph node section (241)

Section 241 underwent the same quality control as remaining sections (see above). Cell-wise mean PanCK/CD45 fluorescent intensities were arcsinh-transformed (cofactor 200) and used to identify local minima (using *ggpmisc*’s *find peaks()* function). These were offset by +/-5% of the corresponding value range for PanCK/CD45, respectively, in order to impose slightly more stringent filtering. Cells identified as PanCK+ and CD45- (c.f. Fig. 4d) were clustered as remaining sections (see above), but using reference profiles infered from the primary tumor only (section 242).

