## Supplementary Material for "Tracing colorectal malignancy transformation from cell to tissue scale"

| **Run** | **Slide** | **Section** | **W x H** (mm) | **#FOVs** | **#Cells** (raw\|fil) | | **%** |
|---|---|---|---|---|---|---|---|
| 1 | 1 | 110 | 8.70 x 17.4 | 383 | 694,553 | 668,061 | 96.2 |
| | 2 | 120 | 11.8 x 15.9 | 396 | 741,155 | 720,282 | 97.2 |
| 2 | 1 | 210 | 7.16 x 11.7 | 212 | 617,498 | 588,750 | 95.3 |
| | 2 | 221 | 9.09 x 6.65 | 136 | 370,074 | 351,968 | 95.1 |
| | 2 | 222 | 6.59 x 5.98 | 73 | 140,542 | 131,095 | 93.3 |
| | 3 | 231 | 4.60 x 12.8 | 146 | 298,151 | 278,691 | 93.5 |
| | 3 | 232 | 4.59 x 5.63 | 71 | 130,814 | 123,600 | 94.5 |
| | 4 | 241* | 5.29 x 2.56 | 33 | 87,976 | 82,692 | 94.0 |
| | 4 | 242 | 12.8 x 8.19 | 188 | 474,471 | 451,327 | 95.1 |
| $\Sigma$ | 6 | 9 | | 1,638 | 3,555,234 | 3,396,466 | 95.5 |

**Table S1: CosMx SMI data overview.** Data were acquired in two runs with two and four slides, respectively, and one or two sections per slide. Width (W) and height (H) correspond to largest difference in tissue-wide x- and y-coordinates, respectively. The scanning area amounts to (512um)$^2$ per field of view (FOV), i.e., 1,638 $\times$ (512um)$^2$ $\approx 4,29$cm$^2$ across all sections. Cell numbers correspond to segmented observations (raw) and those retained after quality control (fil). *Lymph node to section 242

| **Run** | **Section** | **#Cells** (raw\|fil.) | | **%** |
|---|---|---|---|---|
| 1 | 110 | 37,032 | 32,665 | 88.2 |
| | 120 | 33,289 | 30,186 | 90.7 |
| 2 | 210 | 9023 | 8772 | 97.2 |
| | 221 | 2366 | 2303 | 97.3 |
| | 222 | 10,540 | 9980 | 94.7 |
| | 240 | 20,792 | 19,690 | 94.7 |
| 3 | 230 | 12,848 | 12,273 | 95.5 |
| $\Sigma$ | 8 | 125,890 | 115,869 | 92.0 |

**Table S2: snPATHO-seq data overview.**

| section | region | dysplasia/ growth pattern | inflammation severity | inflammation cell type | grading/ staging | fibrous desmoplasia | angio-genesis | necrosis | LI | TLS |
|---|---|---|---|---|---|---|---|---|---|---|
| 110 | REF | | 3 | P | | | | | | |
| | TVA1 | low | 3 | P | | | | | | |
| | TVA2 | low | 3 | M | | | | | | |
| | TVA3 | high, F | 2 | M | | | | | | |
| | CRC | V | 1 | P | G2/T0 | – | – | + | – | + |
| 120 | REF | | 2 | P | | | | | | |
| | TVA | low | 2 | P | | | | | | |
| | CRC1 | A | 4 | P | G1/T1 | + | - | + | – | – |
| | CRC2 | A | 3 | P | G1/T2 | – | + | – | + | + |
| 210 | REF | | 3 | M | | | | | | |
| | TVA | low, F | 2 | M | | | | | | |
| | CRC | M | 4 | P | G1/T2 | – | + | – | – | + |
| 221 | REF | | 2 | M | | | | | | |
| | TVA1 | low | 3 | P | | | | | | |
| | TVA2 | low | 2 | P | | | | | | |
| | TVA3 | low | 3 | P | | | | | | |
| | CRC | A | 4 | P | G1/T2* | + | – | – | + | – |
| 222 | REF | | 4 | P | | | | | | |
| | TVA | low | 3 | P | | | | | | |
| 231 | REF | | 4 | M | | | | | | |
| | TVA | low | 4 | P | | | | | | |
| | CRC | M | 4 | P | G3/T2 | + | + | – | + | + |
| 232 | REF | | 3 | M | | | | | | |
| | CRC | M | 4 | P | G3/T2 | + | – | – | + | + |
| 242 | REF | | 3 | P | | | | | | |
| | TVA | high, F | 4 | P | | | | | | |
| | CRC | A, N | 3 | P | G3/T3 | + | + | + | + | – |

**Table S3: Histopathological metadata.** Rows correspond to regions, grouped by section and colored by domain, i.e., reference-like mucosa (REF), tubulovillous adenoma (TVA), and colorectal carcinoma (CRC); columns correspond to different histopathological classification and characterization criteria: TVA dysplasia (low- or high-grade) and F-focal villous like; and CRC growth pattern (V-villous, A-adenomatous, M-mucinous, N-neuroendocrine); inflammation severity (1-minimal, 2-mild, 3-moderate, 4-high) and cell type (M-mononuclear, P-mixed mononuclear and polymorphonuclear); CRC grading/differentiation (G1-well, G2-moderately, G3-badly differentiated) and CRC staging/invasion (T0-mucosa, T1-submucosa, T2-muscular, T3-serosa); CRC-associated features (LI-lymphovascular invasion, TLS-tertiary lymphoid structure); –/+ = no/yes. *Tissue not well oriented, impairing invasion scoring.

| | 110 | 120 | 210 | 221 | 222 | 231/2 | 241/2 |
|---|---|---|---|---|---|---|---|
| age | 56 | 81 | 56 | 81 | 74 | 70 | 54 |
| gender | M | M | F | F | F | M | M |
| stage | LGD-TVA | pT2(m)N0 | pT2N0 | pT1N0 | – | pT2 | pT1N1b |
| MSS/MSI | x | x | MSS | MSS | – | MSS | MSS |
| CMS | x | x | CMS3 | CMS2 | x | x | x |
| iCMS | x | x | iCMS2 | iCMS2 | x | x | x |
| kras | x | x | Exon 2: c.35G>T (p.(Gly12Val)) | Exon 2: c.38G>A (p.(Gly13Asp)) | – | – | – |
| braf | x | x | WT | WT | – | – | – |
| tp53 | x | x | WT | Exon 5: c.524G>A (p.(Arg175His)) | – | – | – |
| pik3ca | x | x | WT | WT | – | – | – |
| apc | x | x | WT | Exon 15: c.3916G>T (p.(Glu1306*)) | – | – | – |
| fbxw7 | x | x | WT | WT | x | x | x |
| met | x | x | WT | WT | x | x | x |
| stk11 | x | x | WT | WT | x | x | x |
| smad4 | x | x | WT | WT | x | x | x |
| pten | x | x | – | – | x | x | x |
| other | x | x | – | – | x | x | x |

**Table S4: Clinical metadata.** Sections 231/2 and 241/2 come from the same patient each. MSS/MSI = microsatellite (in)stability, (i)CMS = (intrinsic) consensus molecular subtype, LGD = low-grade dysplasia, WT = wild type, – = none/not applicable, x = not available.

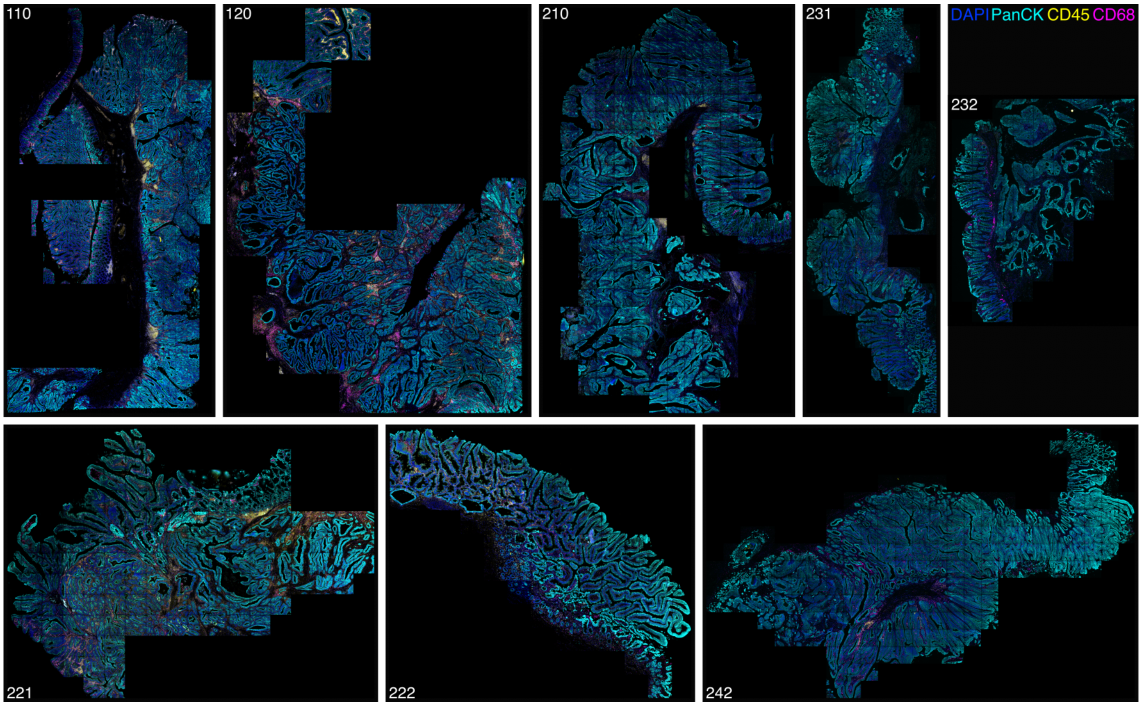


**Fig. S1: Immunofluorescence composite images** including DAPI (nuclei), PanCK (epithelia), CD45 and CD68 (immune). Images are rescaled for better visibility; c.f., Table S1 for factual section dimensions.

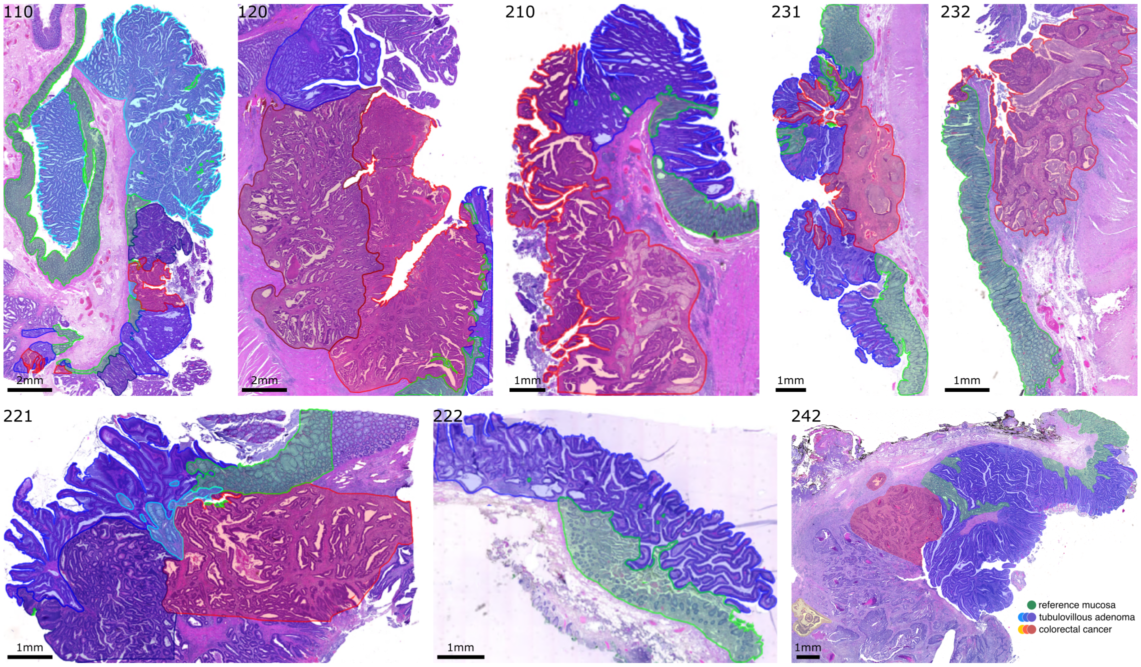


**Fig. S2: Hematoxylin and eosin stains** overlaid with histopathological domains demarcating reference-like mucosa (green), premalignant tubulovillous adenoma (blue), and malignant colorectal cancer (red); regions of the same domain in a section are shown in different color tones.

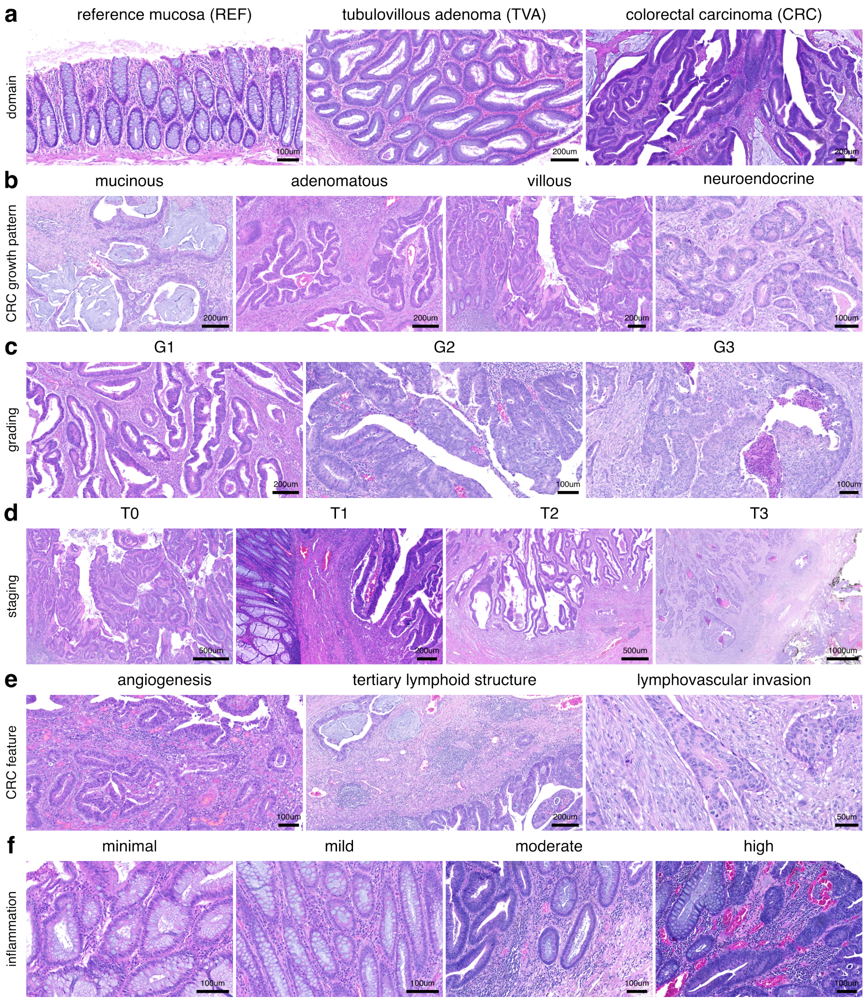


**Fig. S3: Histopathological features and classification criteria.** Shown are representative exemplary H&E stains from across sections. **(a)** Distinct histopathological domains: reference mucosa (REF), premalignant tubulovillous adenoma (TVA) and colorectal carcinoma (CRC). **(b)** Malignant colorectal cancer histological subtypes. **(c)** CRC tumor differentiation. **(d)** CRC tissue invasion depth. **(e)** Relevant cancer-associated features. **(f)** Inflammatory infiltration severity.

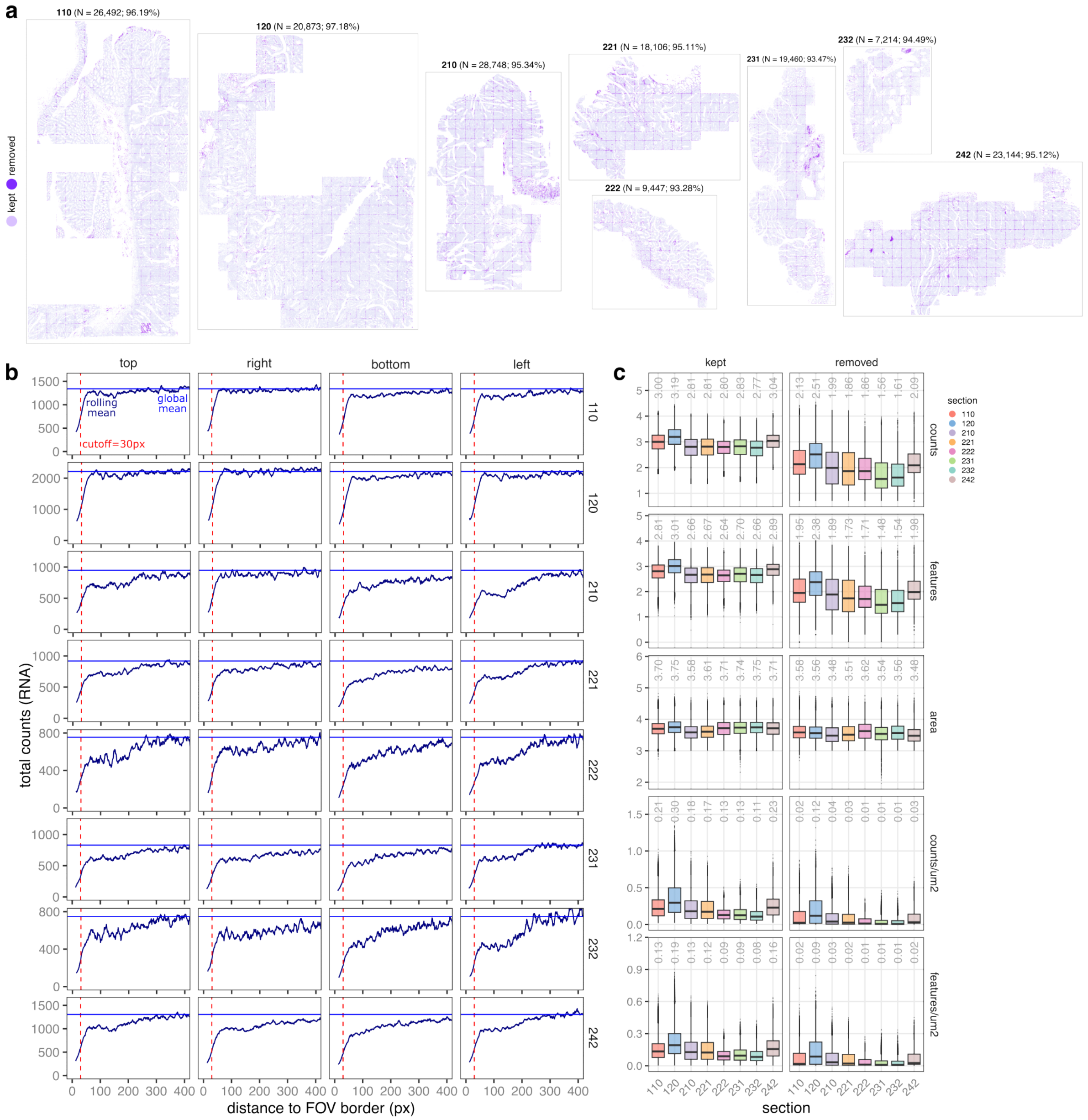


**Fig. S4: Quality control.** **(a)** Spatial plots highlighting low-quality cells. **(b)** Average total RNA counts at a given distance to different field of view (FOV) borders (px); x- and y-axis values correspond to rolling means (window size 10px). **(c)** Boxplots of (fttb) total RNA counts, uniquely detected features, and cell area (um$^2$), as well as total counts and unique features normalized for area; non-normalized y-axis values are $\log_{10}$-transformed, numbers denote medians.

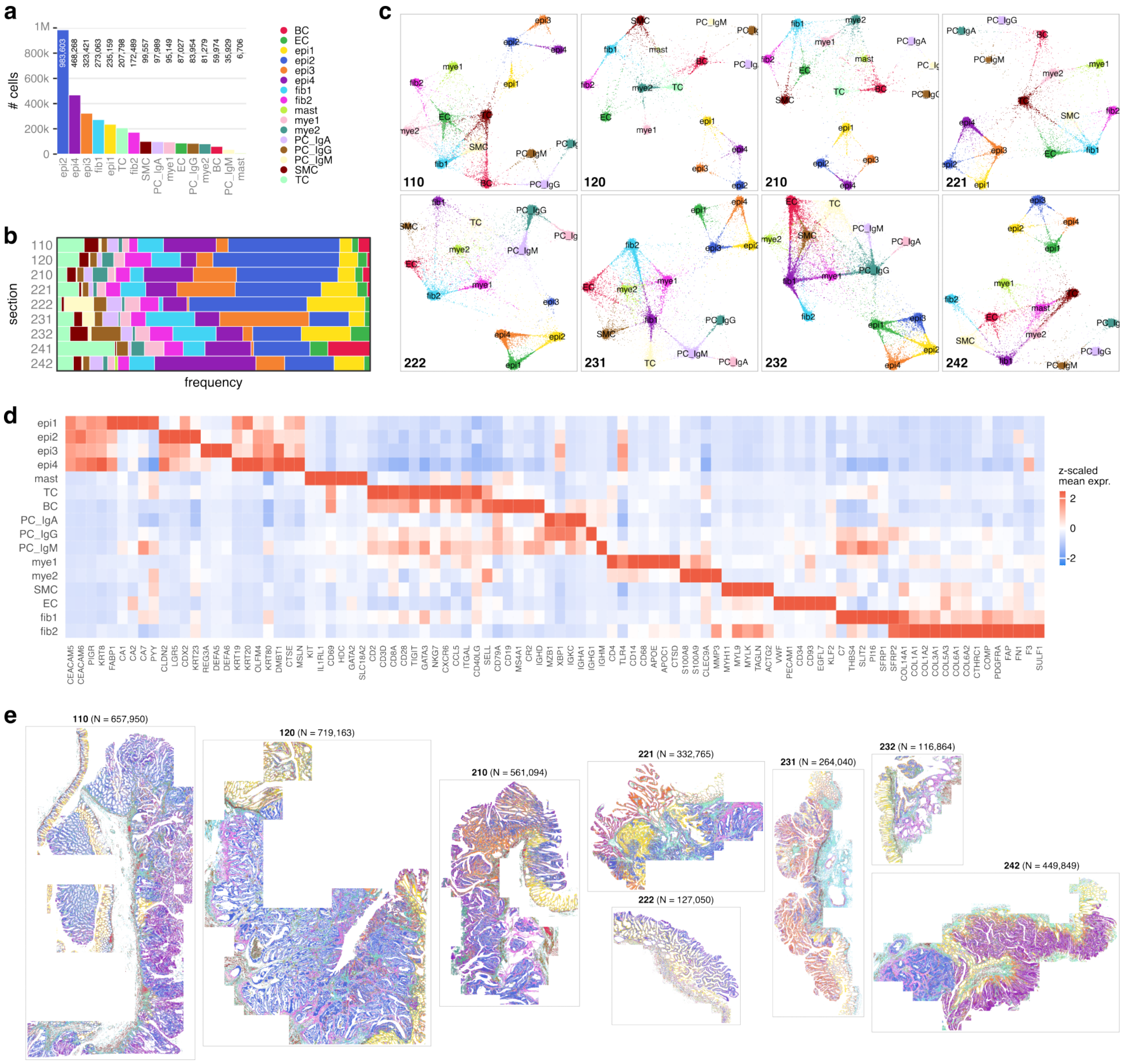


**Fig. S5: Low-resolution clustering results.** **(a)** Number of cells assigned to each subpopulation overall. **(b)** Subpopulation frequencies across sections. **(c)** Section-wise 'flightpath' plots from *InSituType* clustering, i.e., UMAP embeddings of posterior assignment probabilities; points = cells downsampled to at most 10,000 per subpopulation. **(d)** Heatmap of select marker genes (data are across cells from all sections). **(e)** Spatial plots with cells colored by subpopulation.

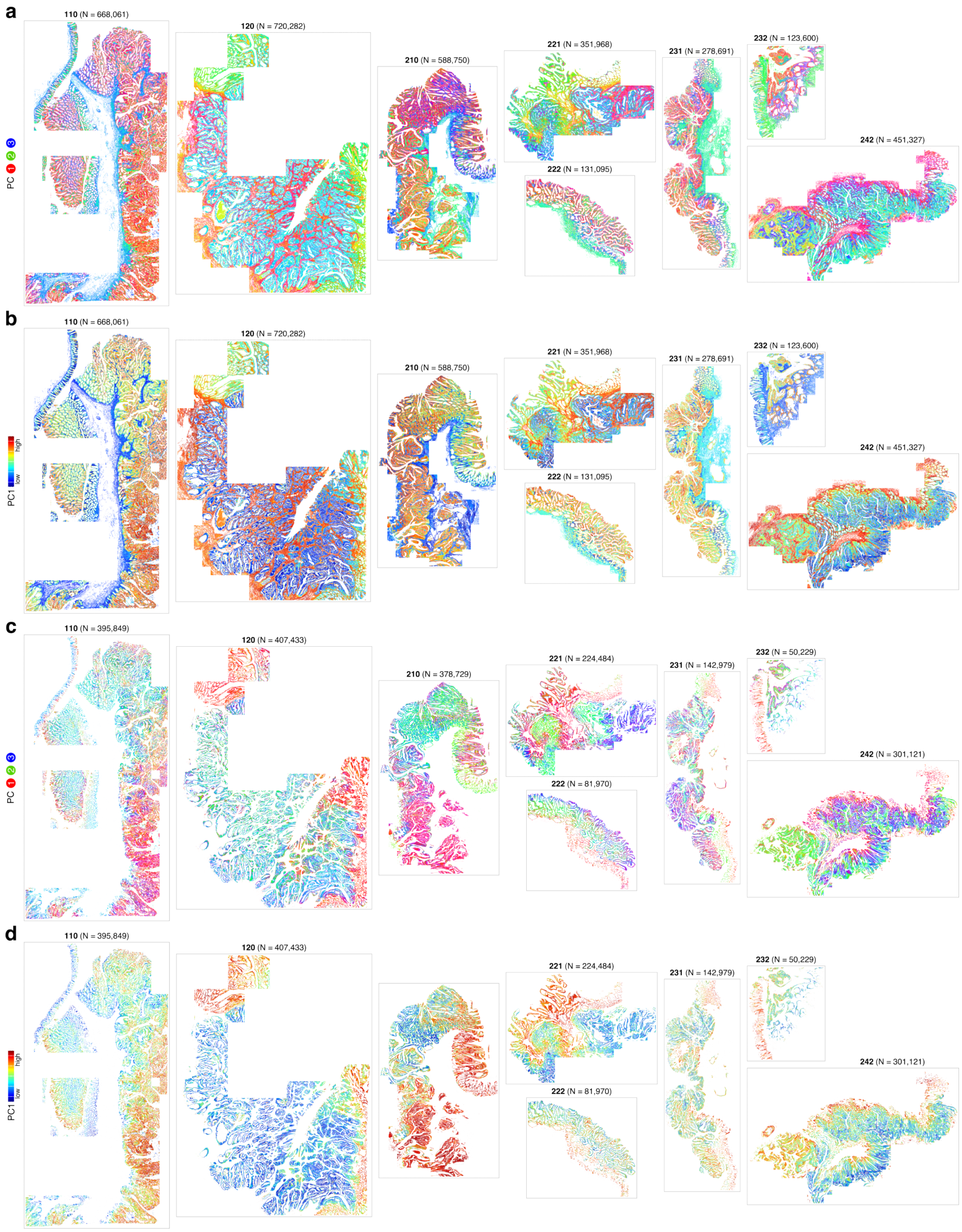


**Fig. S6: Spatial plots of principal components.** Cells are colored by **(a)** PCs 1-3 represented as RGB colors, and **(b)** PC1 values (rescaled using 1 and 99 percentiles as boundaries). **(c-d)** Same plots restricted to epithelial cells.

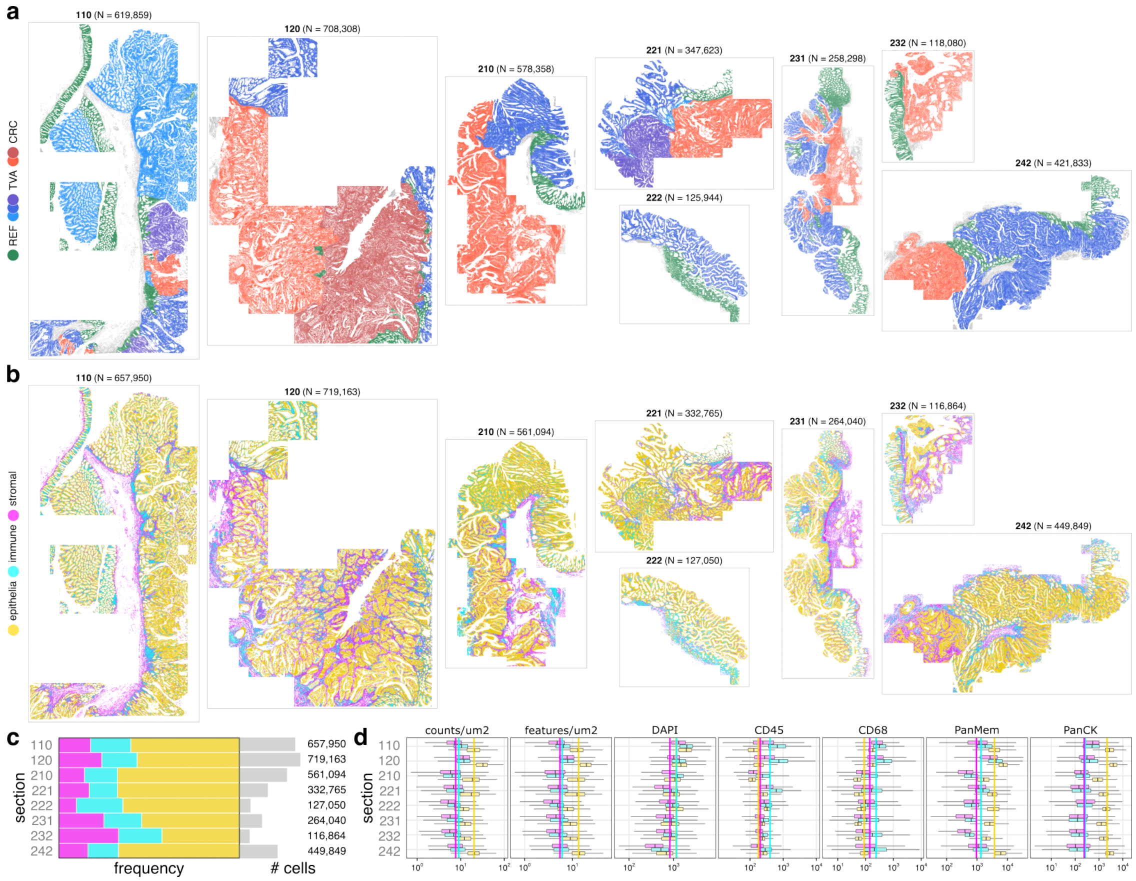


**Fig. S7: Histopathological regions and compartment-level annotations.** **(a)** Spatial plots with cells colored by histopathological classification into regions of reference-like mucosa (REF), premalignant tubulovillous adenoma (TVA), and malignant colorectal cancer (CRC). **(b)** Spatial plots of compartment-level annotations into epithelia, immune, and stromal subpopulations. **(c)** Relative abundance of compartments across sections. **(d)** Total RNA counts, uniquely detected RNA targets, and mean immunofluorescence signal across section, stratified by compartment.

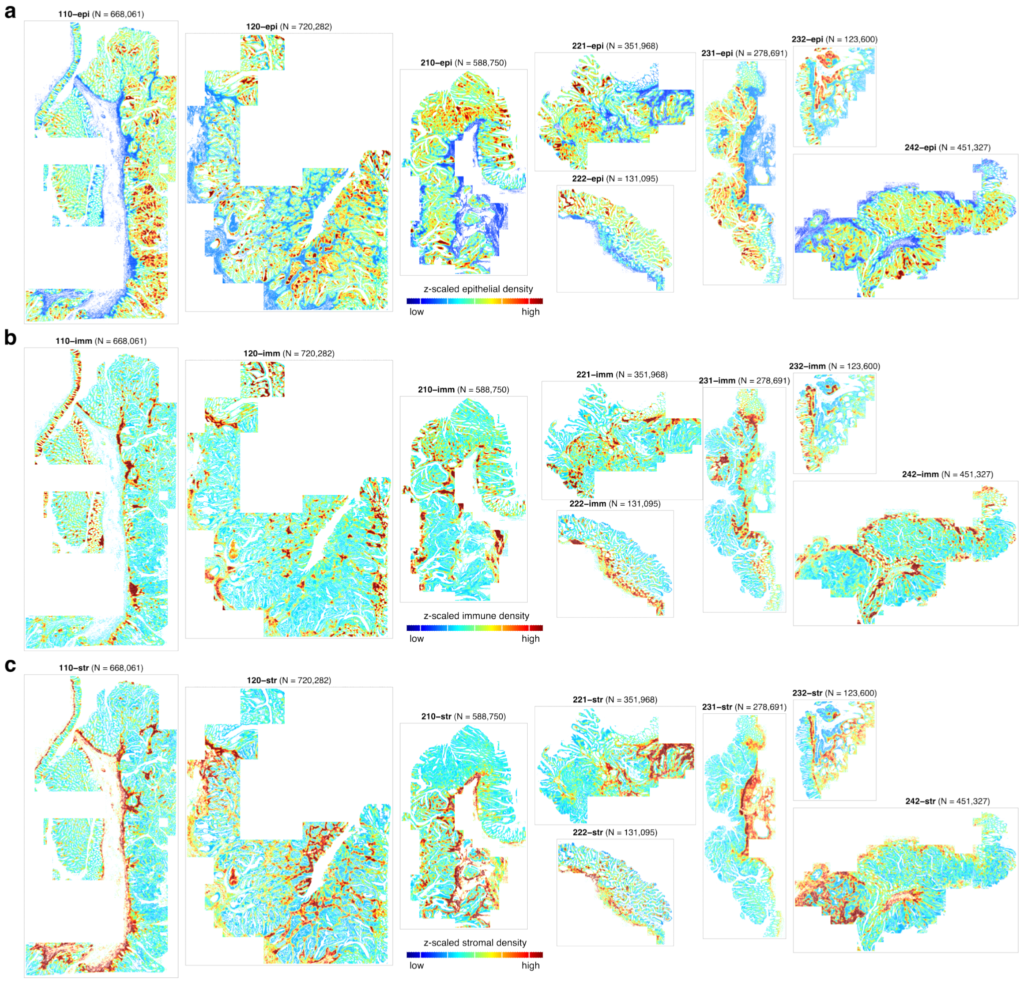


**Fig. S8: Cellular density, stratified by compartment.** Spatial plots with cells colored by the local density of **(a)** epithelial, **(b)** immune, and **(c)** stromal subpopulations, estimated as the number of cells from a given compartment within a 50um radius from each cells, z-scaled across cells in a given section.

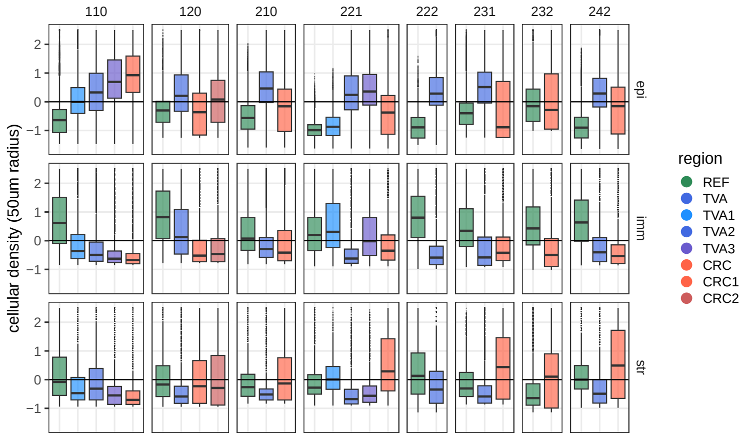


**Fig. S9: Comparison of cellular densities across histopathological regions.** Estimates correspond to the number epi(thelial), imm(une), and str(omal) cells within a 50um radius, z-scaled across cells in a given section (column) and subset (row); i.e., values are comparable within but not across panels. For each section and subset, data are stratified by histopathological regions (c.f. Table S3 and Fig. S2).

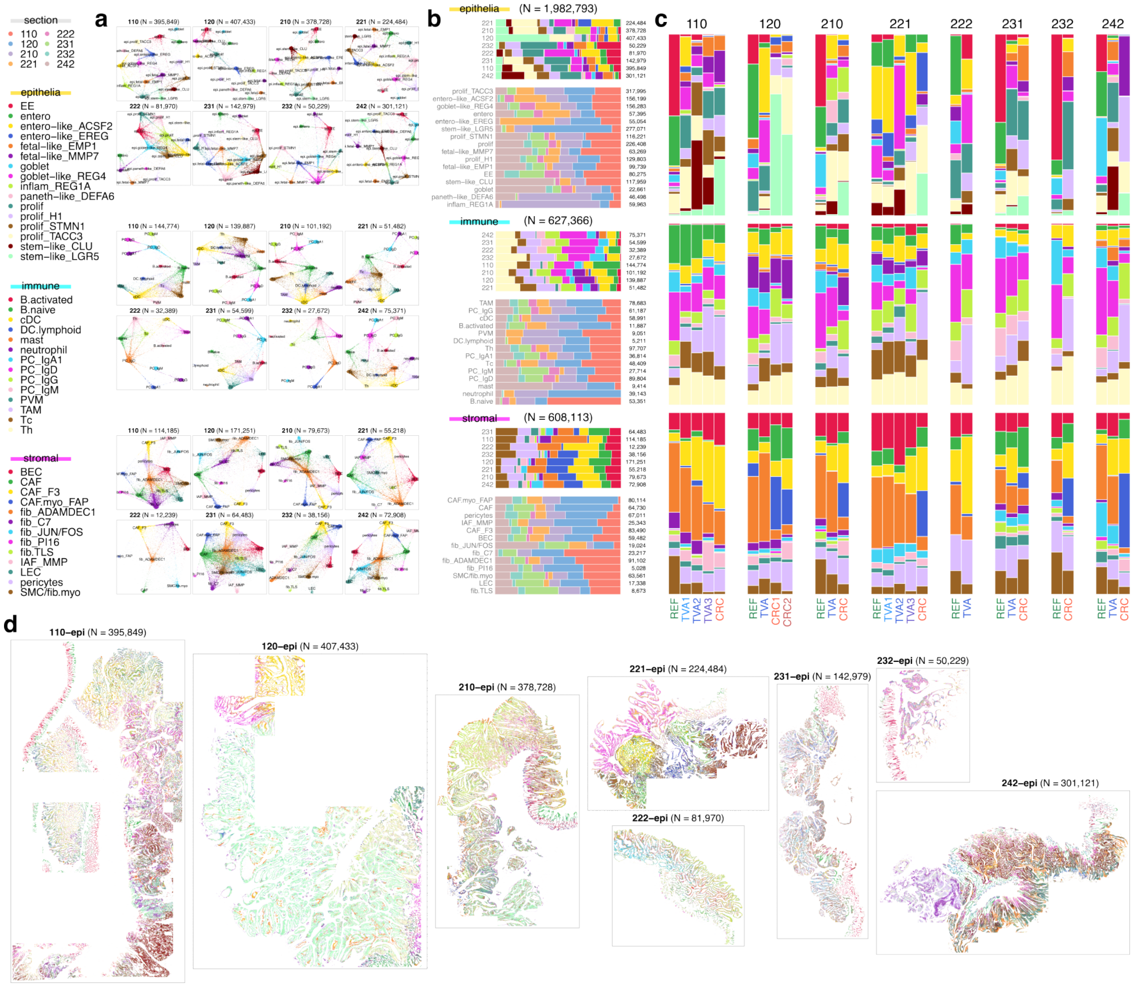

**Fig. S10: Clustering results.** **(a)** Section-wise 'flightpath' plots from *InSituType* clustering, i.e., UMAP embeddings of posterior assignment probabilities; points = cells downsampled to at most 10,000 per subpopulation. **(b)** Subpopulation frequencies across sections; y-axes are ordered by hierarchical clustering. **(c)** Subpopulation frequencies across histopathological regions; rows = compartments, columns = sections, x-axes ordered by malignancy. **(d)** Spatial plots of epithelial cells colored by subpopulation.

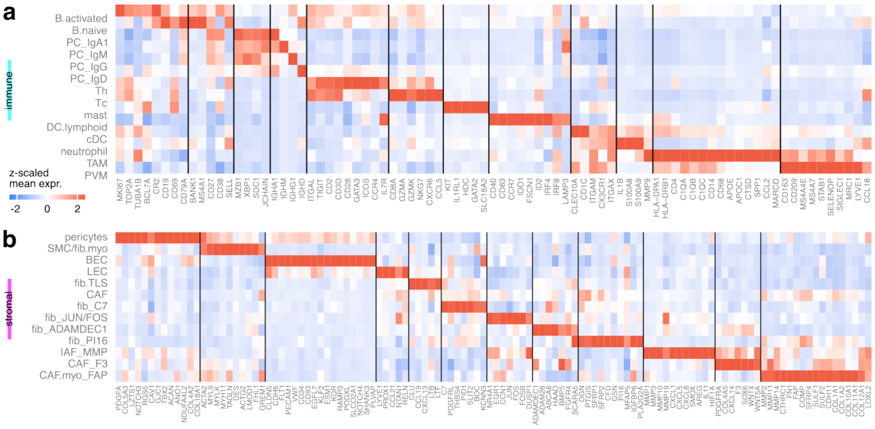

**Fig. S11: Selected marker genes** for (a) immune and (b) stromal subpopulations. Heatmap values correspond to expression values (log-library size normalized counts) averaged across cells from all sections, and z-scaled across clusters (thresholded at $|2.5|$ standard deviations).

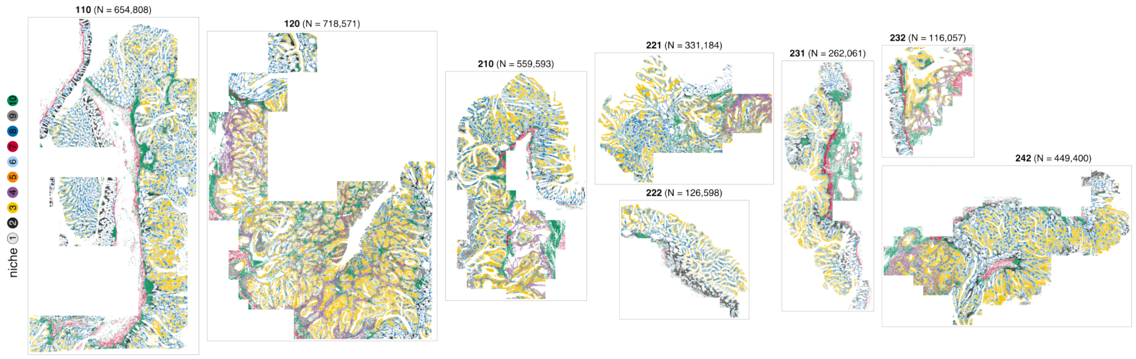


**Fig. S12: Niche analysis.** Spatial plots with cells colored by niche (N1-10). Assignments are based on quantifying subpopulation frequencies (equating epithelial identities) among each cell's 50um radial neighborhood, and using the resulting (cells $\times$ clusters) matrix of proportions as input to $k$-means clustering with $k$=10.

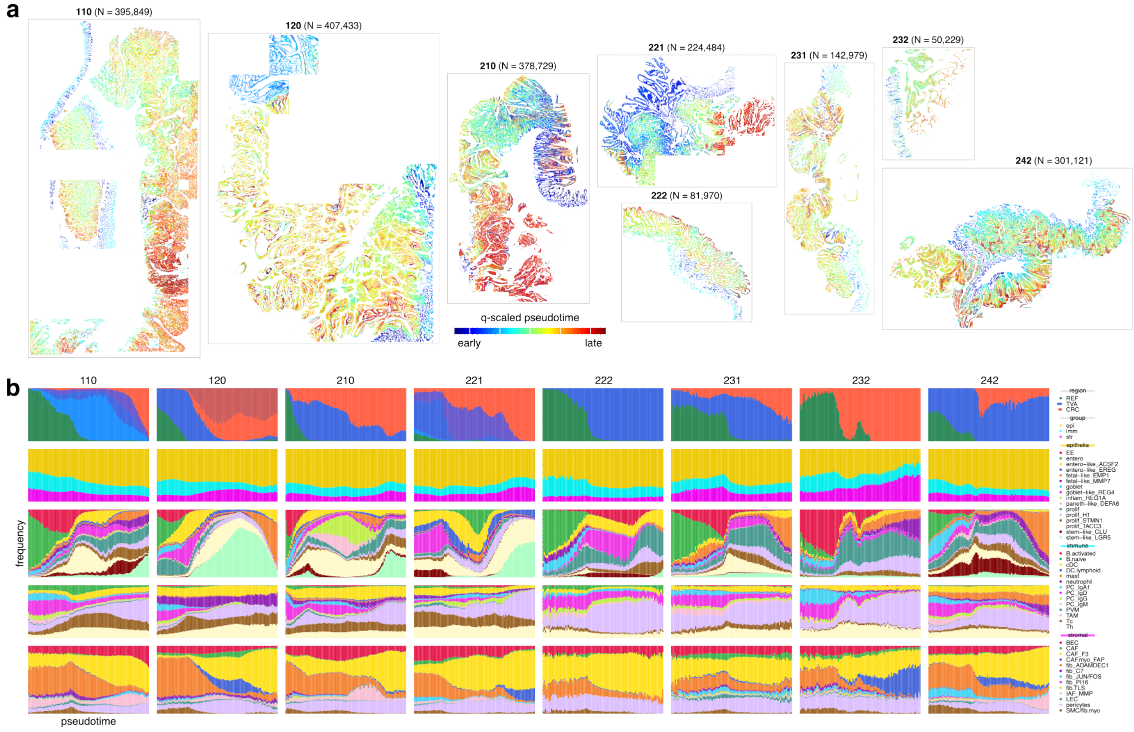


**Fig. S13: Epithelial trajectory inference.** **(a)** Spatial plots with epithelial cells colored pseudotime values (scaled between 0 and 1 using 1%- and 99%-quantiles as boundaries within each section). **(b)** Frequency of histopathological regions (epithelia only), compartment-level and -wise subpopulations over pseudotime; for non-epithelia, values correspond to the proportion of cells within a 50um radial neighborhood.

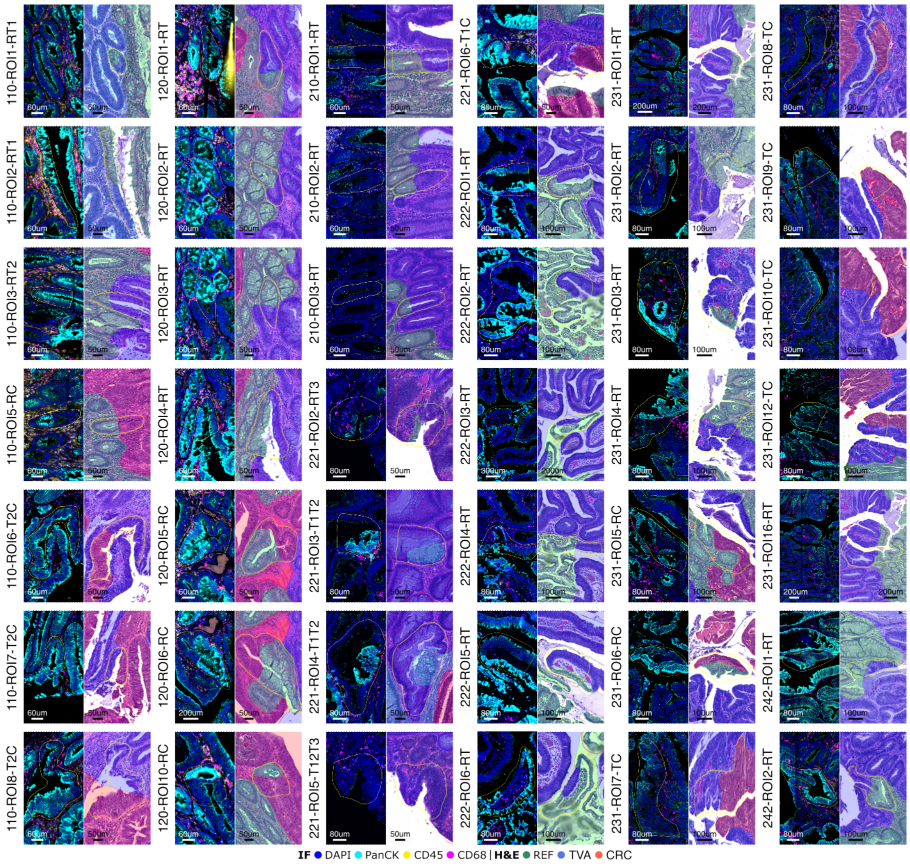


**Fig. S14: Transition crypts** between histopathological domains (REF, TVA, CRC). Labels denote section identifier, region of interest (ROI) enumerator, and the transition type: REF to TVA (RT), REF to CRC (RC), TVA to CRC (TC), and between TVA subdomains (TT). Each panel shows one crypt (N=41 in total); left: immunofluorescence (IF) composite image, right: H&E stain overlaid with annotated domains.

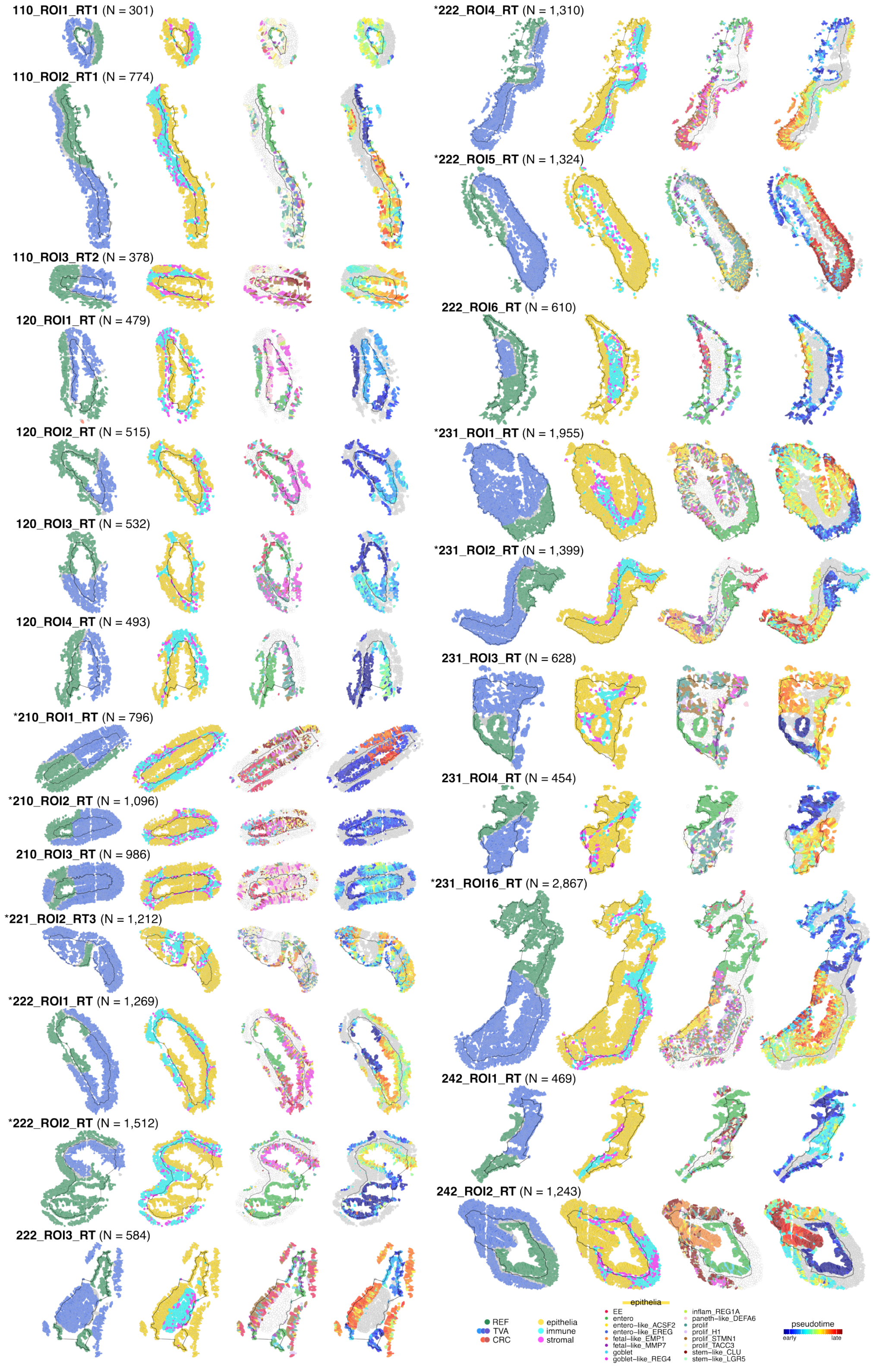


**Fig. S15: Spatial plots of transition crypts** REF-TVA, colored by (fltr) histopathological domain, compartment, epithelial subpopulation, and pseudotime. Black lines pass through centroids of selected cells; all cells within a 50um expanded region are included. *Downsized by a scaling factor of 0.75.

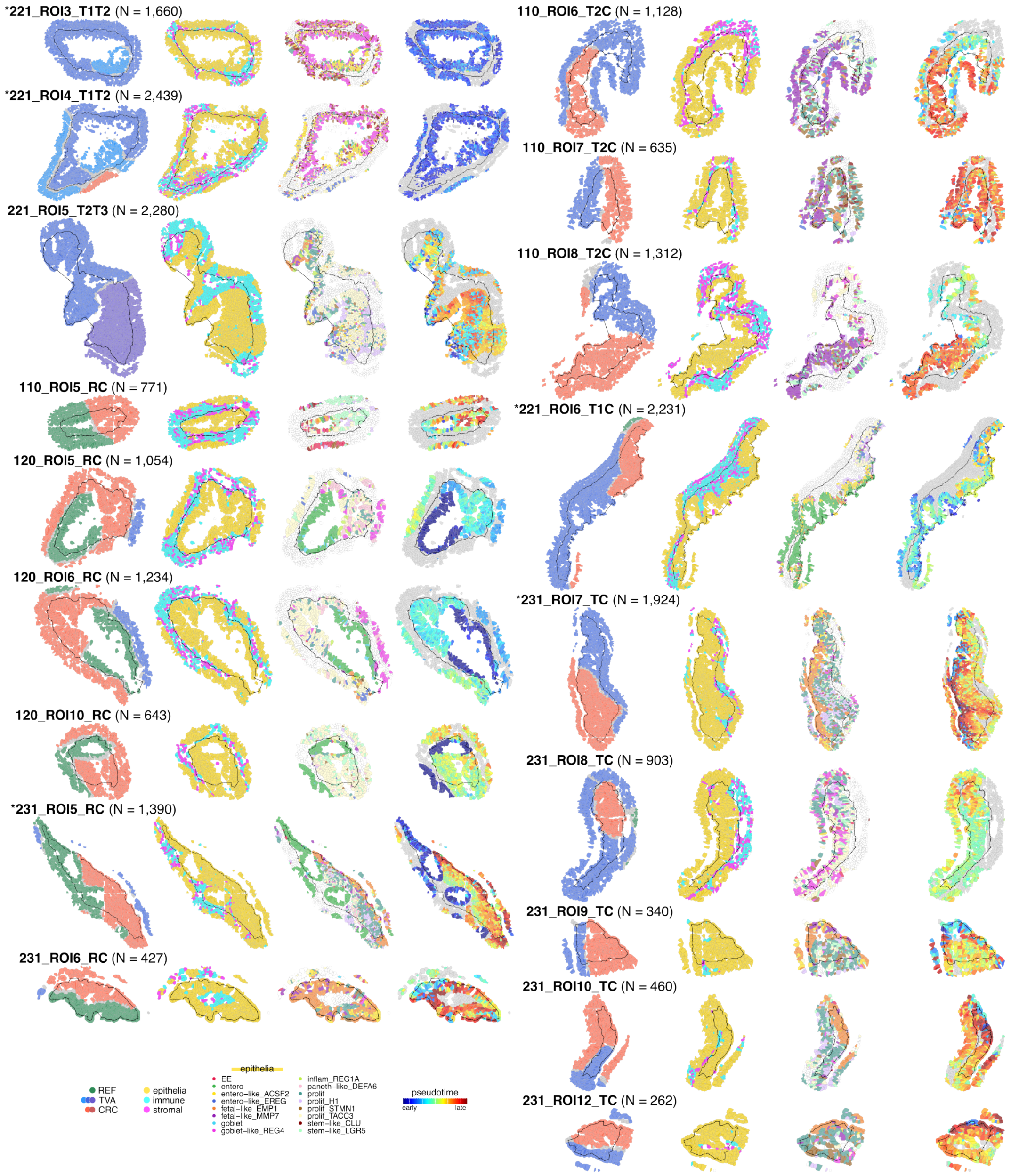


**Fig. S16: Spatial plots of transition crypts** inter-TVA, REF-CRC and TVA-CRC, colored by (fltr) histopathological domain, compartment, epithelial subpopulation, and pseudotime. Black lines pass through centroids of selected cells; all cells within a 50um expanded region are included. *Downsized by a scaling factor of 0.75.

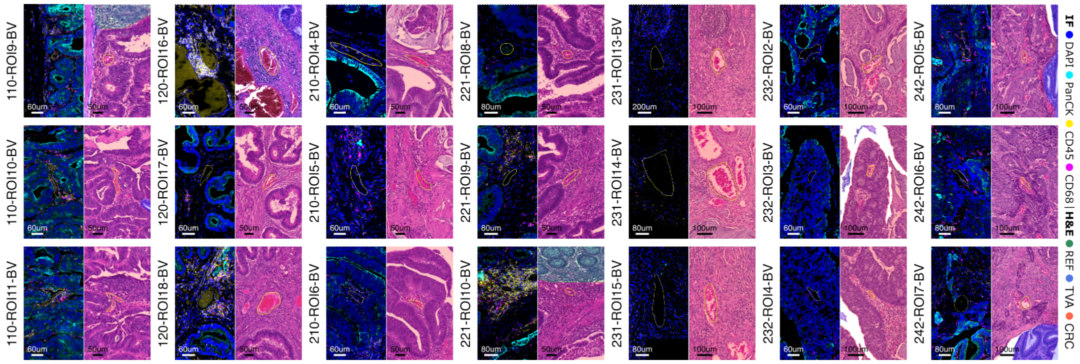

**Fig. S17: Invasion-free blood vessels.** Each panel shows one vessel (3 per section except 222; N=21 in total); left: immunofluorescence (IF) composite image, right: hematoxylin and eosin (H&E) stain overlaid with region annotation (REF = reference-like, TVA = tubulovillous adenoma, CRC = colorectal cancer).

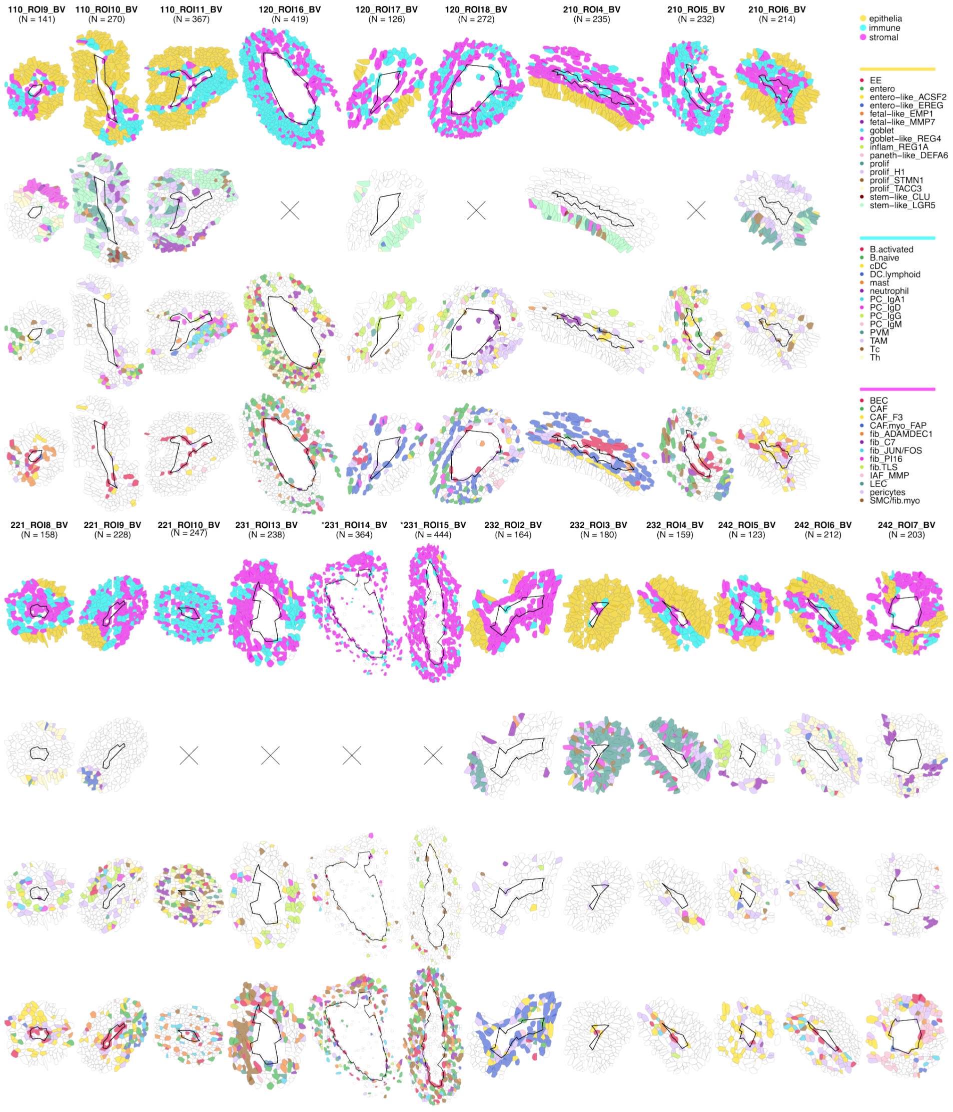

**Fig. S18: Spatial plots of invasion-free blood vessels** colored by (from top to bottom) compartment, epithelial, immune and stromal subpopulation. Black lines pass through centroids of selected cells; all cells within a 50um expanded region are included. Vessels are ordered by section; c.f. Fig. S17. *Downsized by a scaling factor of 0.75; × = placeholder when N≤1.

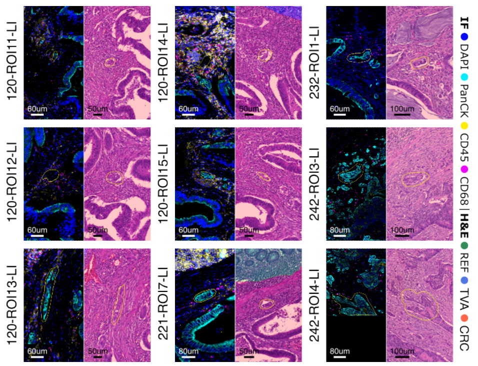


**Fig. S19: Lymphovascular invasions.** Each panel shows one invasions (N=9 in total); left: immunofluorescence (IF) composite image, right: hematoxylin and eosin (H&E) stain overlaid with region annotation (REF = reference-like, TVA = tubulovillous adenoma, CRC = colorectal cancer).

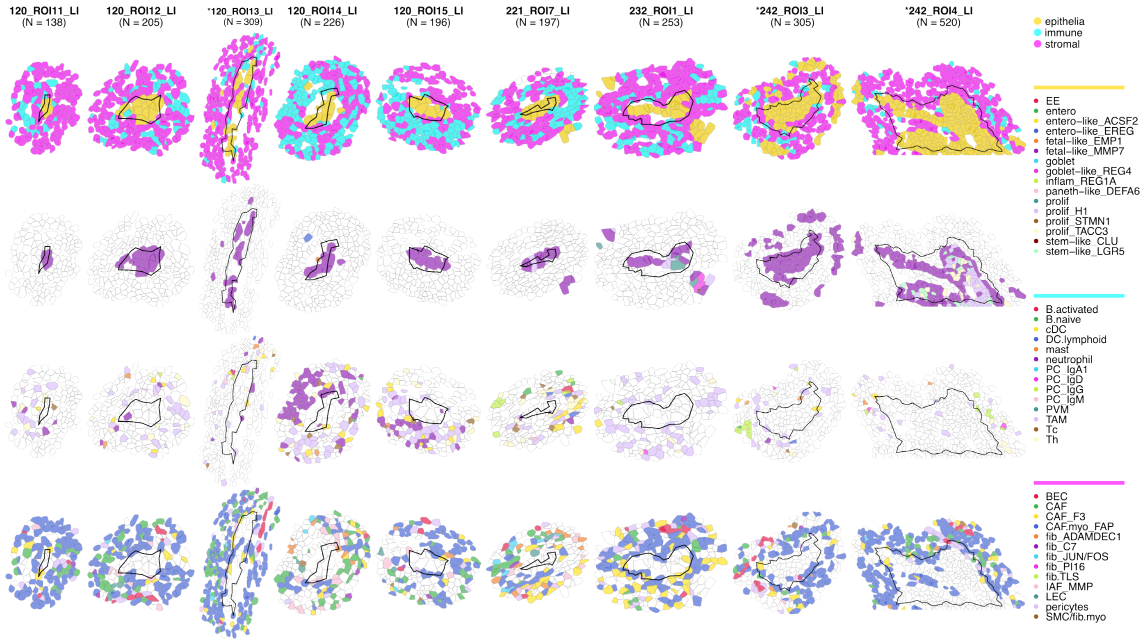


**Fig. S20: Spatial plots of lymphovascular invasions** colored by (from top to bottom) compartment, epithelial, immune and stromal subpopulation. Black lines pass through centroids of selected cells; all cells within a 50um expanded region are included. Invasions are ordered by section; c.f. Fig. S19. *Downsized by a scaling factor of 0.75.
